# Integrated analysis of the methylome and transcriptome of twin almonds (*Prunus dulcis* [Mill.] D.A.Webb) reveals genomic features associated with non-infectious bud failure

**DOI:** 10.1101/2021.02.08.430330

**Authors:** Katherine M. D’Amico-Willman, Chad E. Niederhuth, Matthew R. Willman, Thomas M. Gradziel, Wilburforce Z. Ouma, Tea Meulia, Jonathan Fresnedo-Ramírez

## Abstract

I.

Almond (*Prunus dulcis* [Mill.] D.A.Webb) exhibits an age-related disorder called non-infectious bud-failure (BF) affecting vegetative bud development and nut yield. The underlying cause of BF remains unknown but is hypothesized to be associated with heritable epigenetic mechanisms. To address this disorder and its epigenetic components, we utilized a monozygotic twin study model profiling genome-wide DNA methylation and gene expression in two sets of twin almonds discordant for BF-exhibition. Analysis of DNA methylation patterns show that BF-exhibition and methylation, namely hypomethylation, are not independent phenomena. Transcriptomic data generated from the twin pairs also shows genome-wide differential gene expression associated with BF-exhibition. After identifying differentially methylated regions (DMRs) in each twin pair, a comparison revealed 170 shared DMRs between the two twin pairs. These DMRs and the associated genetic components may play a role in BF-exhibition. A subset of 52 shared DMRs are in close proximity to genes involved in meristem maintenance, cell cycle regulation, and response to heat stress. Annotation of specific genes included involvement in processes like cell wall development, calcium ion signaling, and DNA methylation. Results of this work support the hypothesis that BF-exhibition is associated with hypomethylation in almond, and identified DMRs and differentially expressed genes can serve as potential biomarkers to assess BF-potential in almond germplasm. Our results contribute to an understanding of the contribution of epigenetic disorders in agricultural performance and biological fitness of perennials.

**Significance:** This study examines epigenetic components underlying noninfectious bud failure, an aging-related disorder affecting almond. Results from this work contribute to our understanding of the implications of DNA methylation on agricultural production, namely perennial fruit and nut production, due to effects on growth, development, and reproduction. Describing the methylome of discordant, monozygotic twin almonds enables the study of genomic features underlying noninfectious bud failure in this economically important crop.

## III. Introduction

Plant diseases and disorders are responsible for reductions in performance and fitness of individuals and can be particularly devastating in crop production systems. While plant diseases are typically attributed to pathogens, several disorders of plants have abiotic or non-pathogenic origins (Kennelly *et al.*, 2012). Non-pathogenic disorders in plants can result from issues like nutrient deficiencies or unfavorable climactic conditions (Kennelly *et al.*, 2012), but these disorders can also result from genetic abnormalities leading to genome instability (Tremblay *et al.*, 1999; Henry *et al.*, 2010). Information on the development of (epi)genetic disorders in plants is currently lacking, but there is evidence that genetic abnormalities, including chromatin modifications, can occur with increased plant age and result in undesirable phenotypes (Watson and Riha, 2011; Dubrovina and Kiselev, 2016).

Almond (*Prunus dulcis* [Mill.] D.A.Webb) is an economically-important nut-crop with less than a dozen extensively-grown cultivars and exhibits an age-related disorder called non-infectious bud-failure (BF) (Kester and Jones, 1970; Almond Board of California, 2019). Exhibition of this disorder leads to repression of vegetative meristem development in the spring, indirectly decreasing yield (Wilson and Schein, 1956; Kester and Jones, 1970). BF is transmitted to both vegetative propagules and sexual progeny without a pathogenic origin or epidemiological pattern of dispersion, and the disorder is non-reversible, meaning a tree will always show the phenotype following initial exhibition (Fenton *et al.*, 1988). The severity of exhibition in progeny, though not immediate, is proportional to severity in the parents (Kester *et al.*, 2004). Cultivars and breeding selections exhibiting this disorder show characteristic dieback at the top of the canopy (Fig. 1), and severe levels of BF can be detrimental to productive orchards with up to 50% yield loss in some cultivars (Gradziel *et al.*, 2013). For example, clones of the cultivar ‘Nonpareil’, representing ~40% of U.S. almond production (Almond Board of California, 2019), exhibit BF, making this disorder a threat to the U.S. agricultural economy. Further, BF led to abandonment of promising almond cultivars including ‘Jordanolo’ (Wilson and Schein, 1956). BF-exhibition has also been observed in other almond producing countries including Australia (M.G. Wirthensohn, pers. comm.), Iran (B. Shiran, pers. comm), and Spain (P.J. Martínez-García, pers. comm.). Our current understanding of BF is limited; to date, no biomarkers to screen for early onset and prevent this disorder have been identified or described.

**Figure 1.**
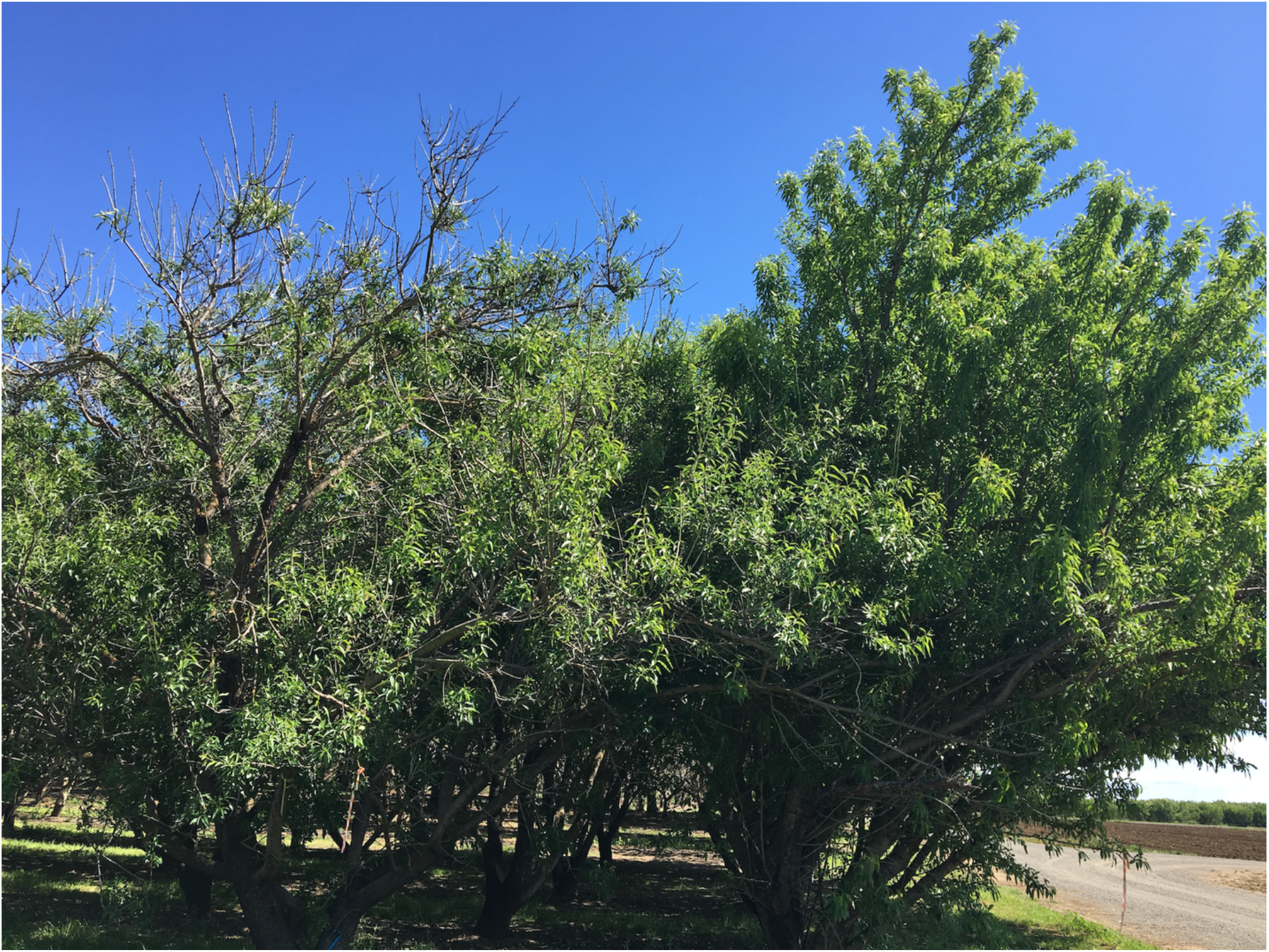
Monozygotic twin almond ‘Stukey’ trees discordant for BF-exhibition; BF twin (left) with typical bud failure signs such as canopy dieback and no-BF twin with common apical growth (right). (Photo taken April 19, 2018 by K. M. D’Amico-Willman)

Given the sexual and asexual transmission along with the environmental cues associated with BF, epigenetic mechanisms may play a role in its exhibition in almond (Kester *et al.*, 2004). DNA methylation is the most widely studied chromatin mark in plants and has been implicated in phenotypic variation in several species including almond (He *et al.*, 2011; Fernández i Martí *et al.*, 2014; Elhamamsy, 2016; Fresnedo-Ramírez *et al.*, 2017). Cytosine methylation occurs in three contexts in plants: CG, CHG, CHH (H = A, T, or G) and can impact gene expression due to inhibition of transcription factors or transcription machinery binding (Zhang *et al.*, 2010; He *et al.*, 2011). Alterations in DNA methylation have been shown to be associated with changes in plant development including meristematic tissue and can be induced as a response to biotic or abiotic stress (Zhang *et al.*, 2010; Bej and Basak, 2017; Seymour and Becker, 2017; Dolzblasz *et al.*, 2018; Alonso *et al.*, 2019). Changes in plant DNA methylation can also be inherited (Richards, 2006; Holeski *et al.*, 2012; Li *et al.*, 2014; Tricker, 2015; Köhler and Springer, 2017), though inheritance of epialleles may not follow the same patterns found in other heritable genetic components (Kakutani, 2002).

Discordant monozygotic (MZ) twin-based studies are an effective tool to precisely address, describe, and quantify DNA methylation mechanisms affecting exhibition of traits within the same genotype. Such studies have been widely used in humans and livestock to identify chromatin marks associated with various disorders (Field and Suttle, 1979; Petronis, 2006; Bell and Spector, 2011; Svendsen *et al.*, 2016). To our knowledge, this framework has not yet been applied in plants despite its potential and suitability in several outcrossing, perennial systems. Almond is an obligated outcrosser (Ushijima *et al.*, 2003) and recalcitrant to inbreeding (Alonso Segura and Socias i Company, 2007; Martínez-García *et al.*, 2012); however, almond can produce fruit with multiple embryos due to post-fertilization embryo splitting resulting in MZ twin almond genotypes (Martínez-Gómez *et al.*, 2002; Martínez-Gómez and Gradziel, 2003). Post-fertilization embryo splitting, often called a double-seed trait, is well known and characterized in the cultivar ‘Nonpareil’ (Martínez-Gómez *et al.*, 2002; Martínez-Gómez and Gradziel, 2003). Currently, almond germplasm derived from MZ twin pairs, known commonly as the ‘Stukey’ collection, is available at the University of California, Davis (Martínez-Gómez *et al.*, 2002), and several twin pairs now exhibit BF. Twin pairs discordant for BF-exhibition in this collection allow for the implementation of a MZ twin-based study model to analyze potential chromatin alterations associated with the disorder.

Fresnedo-Ramírez *et al.* (2017) postulated that DNA methylation interacts with genotype and chronological age in BF-exhibition in almond. However, DNA methylation signatures identified in the prior study were anonymous and qualitative (e.g. presence vs. absence of bands), limiting a precise contrast and characterization of the features influenced by methylation and the role they might play in BF development. Differential DNA methylation signatures associated with BF-status between isogenic individuals may suggest a methylome role in BF-exhibition. Thus, identifying and quantifying these signatures utilizing a MZ twin-based design and an annotated genome will enhance elucidation of the role of DNA methylation in BF-exhibition in almond. In this study, we tested this hypothesis by profiling DNA methylation and gene expression patterns in two sets of MZ twin almonds with discordant BF-exhibition. To our knowledge, this is the first study performed in an outcrossing plant species utilizing a twin-based model and a bisulfite sequencing approach to address an age-related disorder. The DNA methylation profiles generated will aide in further interrogation and subsequent design of strategies to monitor and mitigate BF in almond germplasm.

## IV. Results

### Contrasting genome-wide DNA methylation status in the ‘Stukey’ twins

Whole-genome bisulfite sequencing was performed on four ‘Stukey’ individuals representing two MZ twin pairs discordant for BF-exhibition. Twin 1a exhibits BF while twin 1b is BF-free (i.e. no-BF), and in twin pair two, twin 2a exhibits BF while twin 2b is BF-free. Genome-wide sequencing depth ranged from 70-110X for each individual, and mapping efficiencies following alignment to the ‘Nonpareil’ reference genome ranged 36.5-39.1% representing 25-43X genome-wide coverage (Table S1). Non-methylated cytosine conversion efficiencies were >98% for each individual.

CG methylation is the most abundant genome-wide methylation context for all twins, followed by CHG and CHH, respectively (Table S2). Percent methylation calculations show that in both twin pairs, methylation is higher in the no-BF twin for CHG and CHH contexts (Table S2). A lack of independence between BF-exhibition (represented as a binomial presence/absence) and DNA methylation in each methylation context was demonstrated by Chi-squared (χ^2^) analysis with Yate’s continuity correction (Table S3a-c; p-value < 2.2×10^−16^). While results showed a lack of independence between methylation and BF-status in the CG context, there is an opposite relationship in twin pair 1 compared to twin pair 2 (Table S3a). Genome-wide cytosine methylation in each context is shown across each of the eight scaffolds of the current ‘Nonpareil’ genome assembly (representing the eight chromosomes of the almond genome) with evident regions of higher overall methylation (Fig. S1, Fig. S2, Fig. S3).

### Identifying regions of differential methylation associated with bud failure status

Differentially methylated regions (DMRs) were identified and compared between the BF and no-BF twins in both twin pair one and twin pair two. Significant DMRs were identified in both twin pairs based on percent cytosine methylation comparisons in the three methylation contexts in the respective regions (p-value < 0.01) (Fig. 2). The DMRs identified in each twin pair methylation context combination were further classified based on their proximity to a gene. The DMRs were classified as either within 2,000 base pairs upstream, within 2,000 base pairs downstream, or intragenic (Table 1). Results of the two-tailed permutation test in the CG context show the frequency of DMRs in the upstream and downstream proximity class are significantly higher and the frequency of DMRs in the intragenic class is significantly lower than in null DMR sets randomly distributed throughout the almond genome (Table 1). Permutation testing shows that significantly more DMRs occur in the upstream class for both the CHG and CHH context, significantly fewer DMRs occur in the intragenic class for both contexts, and the frequency of downstream DMRs in both contexts do not exhibit significant deviance from a random frequency distribution of DMRs along the genome (Table 1).

**Table 1.** Number of DMRs in each methylation context for each ‘Stukey’ twin pair. DMRs are classified by proximity relative to a gene: upstream (within 2,000 bp) of a gene, intragenic, or downstream (within 2,000 bp) of a gene (significant permutation tests are represented in bold).

| **Methylation Context** | **Twin pair** | **Proximity** | **Number of DMRs** | **Permutation Result** | **p-value** |
| --- | --- | --- | --- | --- | --- |
| CG | 1 | Upstream | 441 | more | < **0.001** |
|  |  | Intragenic | 154 | less | < **0.001** |
|  |  | Downstream | 339 | more | < **0.001** |
|  | 2 | Upstream | 452 | more | < **0.001** |
|  |  | Intragenic | 175 | less | < **0.001** |
|  |  | Downstream | 368 | more | < **0.001** |
| CHG | 1 | Upstream | 278 | more | < **0.001** |
|  |  | Intragenic | 41 | less | < **0.001** |
|  |  | Downstream | 191 | more | 0.086 |
|  | 2 | Upstream | 328 | more | < **0.001** |
|  |  | Intragenic | 79 | less | < **0.001** |
|  |  | Downstream | 219 | more | 0.427 |
| CHH | 1 | Upstream | 166 | more | < **0.001** |
|  |  | Intragenic | 23 | less | < **0.001** |
|  |  | Downstream | 104 | more | 0.258 |
|  | 2 | Upstream | 229 | more | < **0.001** |
|  |  | Intragenic | 52 | less | < **0.001** |
|  |  | Downstream | 150 | more | 0.200 |

**Figure 2.**
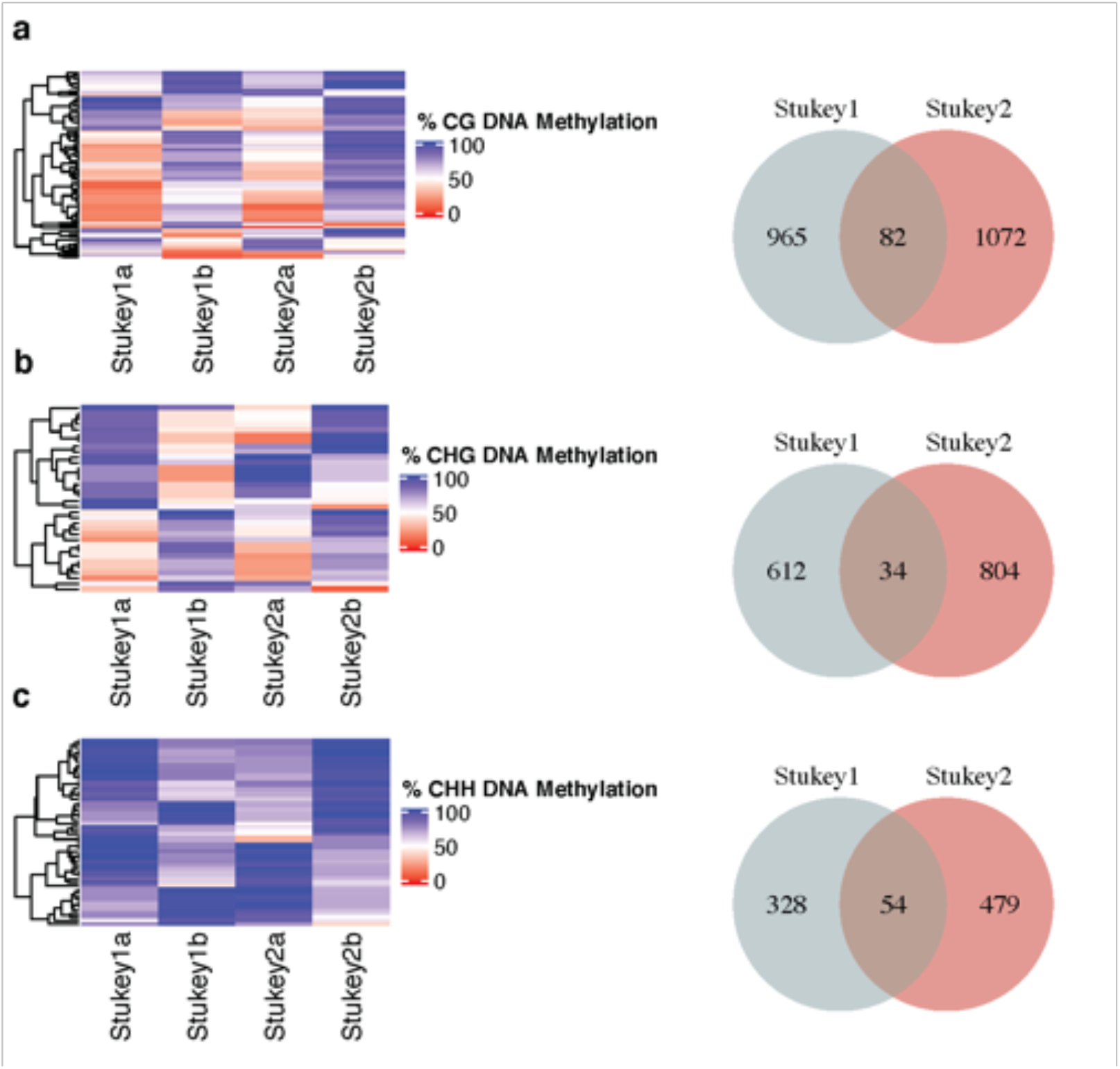
Heatmap displaying percent methylated cytosines in each twin pair for the DMRs in each methylation context: (a) CG methylation, (b) CHG methylation, and (c) CHH methylation. ‘Stukey’ twins 1a and 2a exhibit BF while ‘Stukey’ twins 1b and 2b are BF-free. The Venn diagram represents the total number of significant DMRs in both ‘Stukey’ twin pairs as well as the number of regions shared between the pairs. Panel **(a)** represents the CG context, panel **(b)** represents the CHG context, and panel **(c)** represents the CHH context.

Of the 1,007 significant DMRs identified in the CG context in twin pair one and the 1,154 significant DMRs identified in the CG context in twin pair two, 82 DMRs were associated with the same gene in the same proximity class (Data S1, Fig. 2a). Of the 646 significant DMRs in the CHG context in twin pair one and the 838 significant DMRs in the CHG context in twin pair two, 34 were associated with the same gene in the same proximity class (Data S1, Fig. **2b**). And of the 382 significant DMRs in the CHH context in twin pair one and the 533 significant DMRs in the CHH context in twin pair two, 54 were associated with the same gene in the same proximity class (Data S1, Fig. **2c**). Patterns of percent methylation within each gene associated-DMR are similar for BF twins and no-BF twins (Fig. **2a,b,c**). The DMRs range in length from 51 – 1,460 bp, the average length of the DMRs in all contexts is ~173 bp, the median DMR length is 128 bp, and the most frequent DMR length is 77 bp (Fig. S4).

The genomic coordinates of the DMRs associated with the same gene from each twin pair were compared, and 121 of the 170 DMRs had overlapping genomic coordinates. The DMRs associated with the same gene and proximity class were further classified as either hyper- or hypomethylated in the BF twins compared to the no-BF twins in each methylation context and proximity class. Of these 170 DMRs, 29 were hypermethylated and 68 were hypomethylated. The remaining 73 DMRs were classified as either hyper- or hypomethylated in one twin pair and had the opposite classification in the other twin pair. Comparing percent methylation difference by methylation context in hyper- and hypomethylated-DMRs associated with the same gene shows that most DMRs in the CG context are hypermethylated in BF twins, while DMRs in the CHG and CHH contexts tend to be hypomethylated in BF twins (Fig. **S5a,b,c**). Comparing percent methylation by proximity class for the same set of DMRs shows that all intragenic DMRs are hypermethylated in BF twins, while DMRs upstream and downstream are both hyper- and hypomethylated in BF twins (Fig. **S6a,b,c**). Permutation testing revealed significantly more shared DMRs in all methylation contexts and proximity classes with the exception of intragenic CHG DMRs when comparing to null DMR sets (Table 2).

**Table 2.** Number of identified DMRs per twin pair that are associated with the same gene, are in the same proximity class relative to that gene, and are in the same context in both ‘Stukey’ twin pair 1 and ‘Stukey’ twin pair 2 (significant permutation tests are represented in bold). Values in parentheses represent the number of DMRs from each twin pair that have overlapping genomic coordinates.

| **Methylation Context** | **Proximity** | **Number of DMRs Per Twin Pair** | **Permutation Result** | **p-value** |
| --- | --- | --- | --- | --- |
| CG | Upstream | 42 (36) | more | **< 0.001** |
|  | Intragenic | 10 (6) | more | **0.006** |
|  | Downstream | 30 (24) | more | **< 0.001** |
| CHG | Upstream | 18 (12) | more | **< 0.001** |
|  | Intragenic | 2 (1) | more | 0.179 |
|  | Downstream | 14 (7) | more | **< 0.001** |
| CHH | Upstream | 34 (22) | more | **< 0.001** |
|  | Intragenic | 3 (2) | more | **< 0.001** |
|  | Downstream | 17 (11) | more | **< 0.001** |

The magnitude of methylation difference in each DMR associated with the same gene and proximity class showed a significant linear relationship when comparing percent methylation for each DMR in no-BF twin 1b to the corresponding DMR in no-BF twin 2b in the upstream and downstream proximity classes (Fig. **S7a** – upstream, Fig. **S7c** – intragenic, Fig. **S7e** – downstream). A significant linear relationship was also shown when comparing percent methylation for each DMR in BF twin 1a to the corresponding DMR in BF twin 2a for each proximity class (Fig. **S7b** – upstream, Fig. **S7d** – intragenic, Fig. **S7f** – downstream).

### Annotation of DMR-associated genes identified in the ‘Stukey’ twins

The genomic sequence of 15 genes associated with hypermethylated DMRs significantly aligned to previously identified genes in the UniProtKB database (Table 3). And the genomic sequence of 37 genes associated with hypomethylated DMRs significantly aligned to previously identified genes (Table 3). Gene ontology (GO) enrichment analysis revealed significant enrichment in biological process GO terms: regulation of timing from vegetative to reproductive phase, protein sumoylation, and glucosinate metabolic processes (Table S4a). Enrichment analysis also revealed enrichment of the molecular function GO term, protein tag (Table S4b). GO terms assigned to all DMR-associated genes are also reported (Table S4a, S4b, S4c).

**Table 3.**
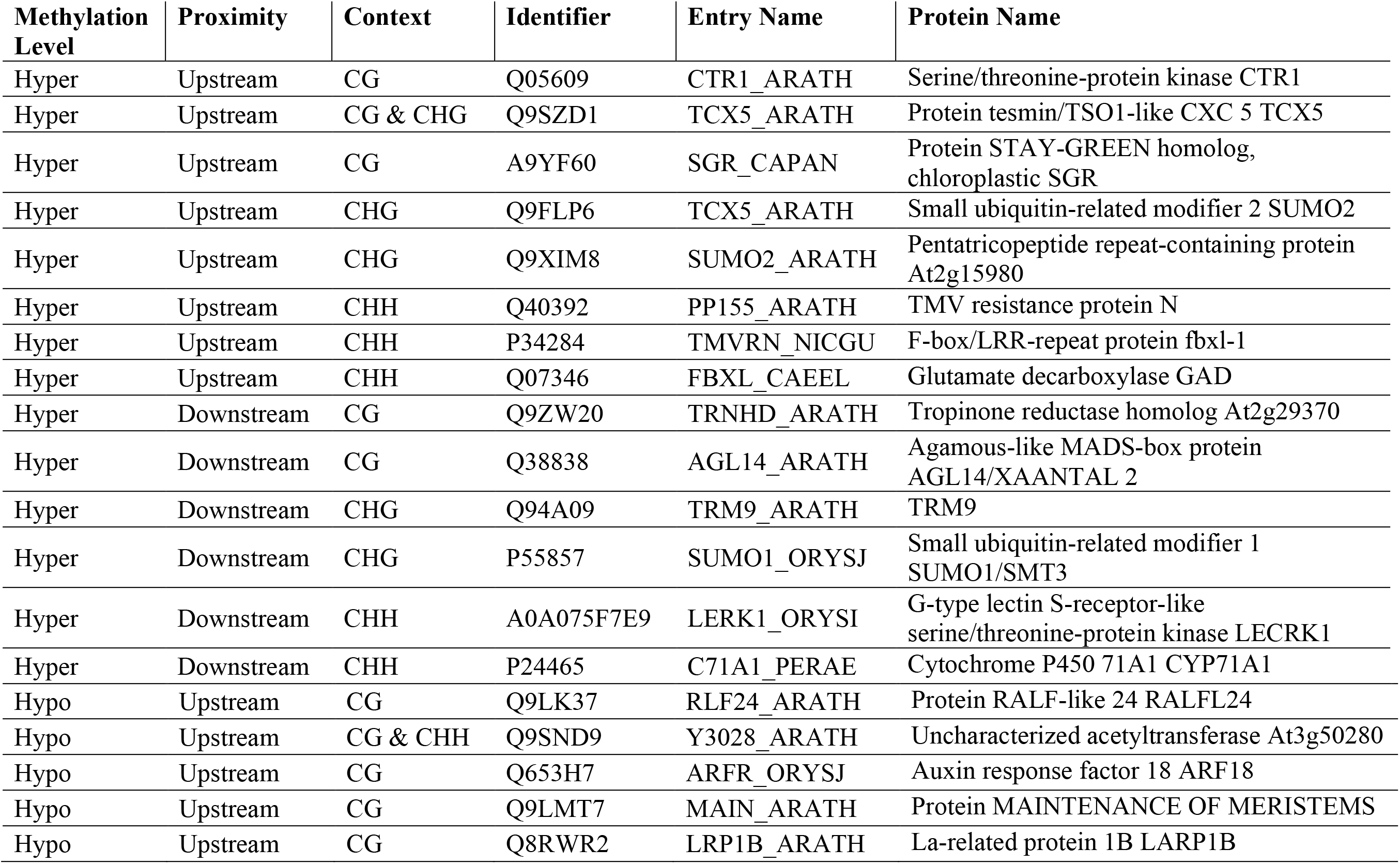
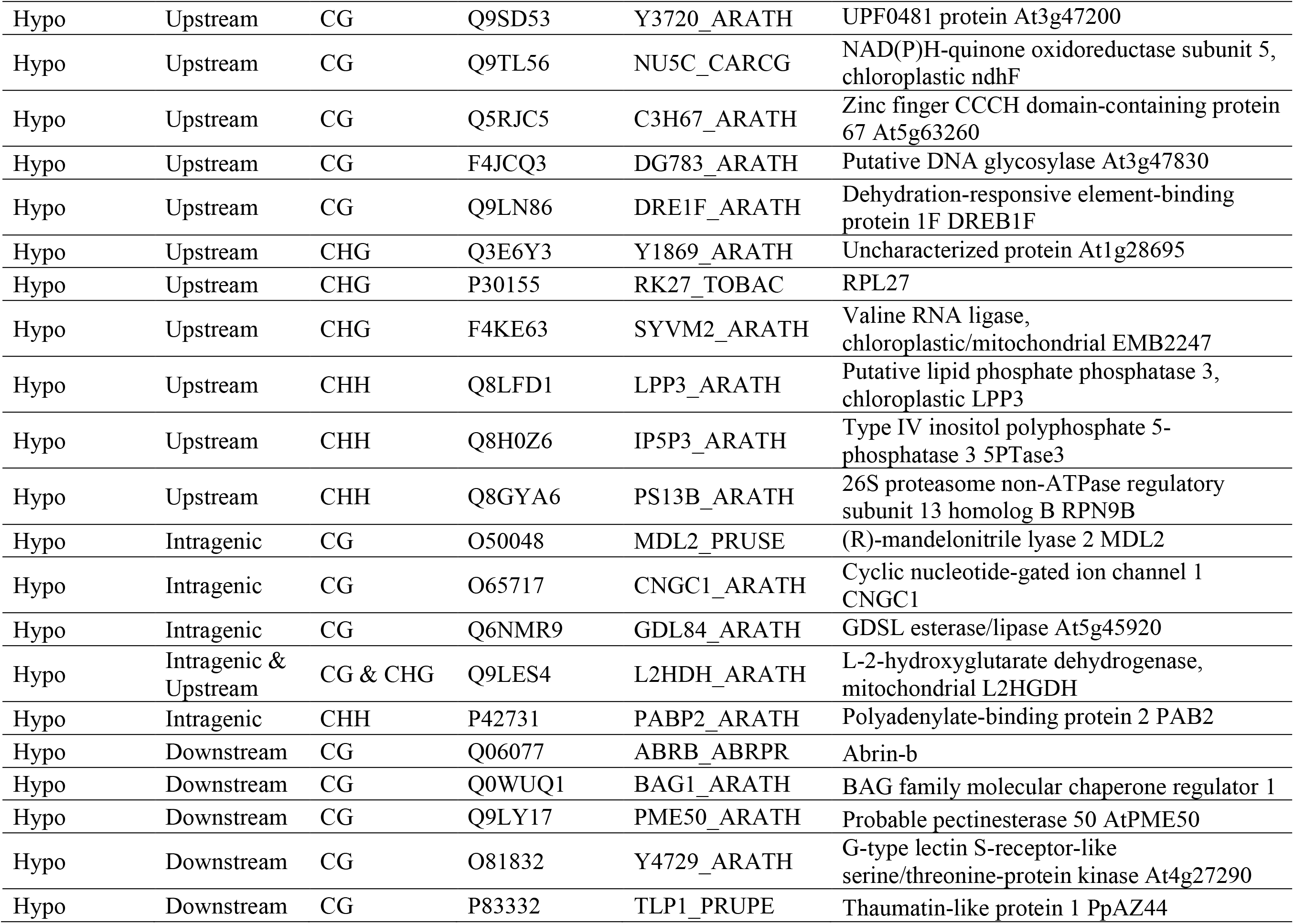
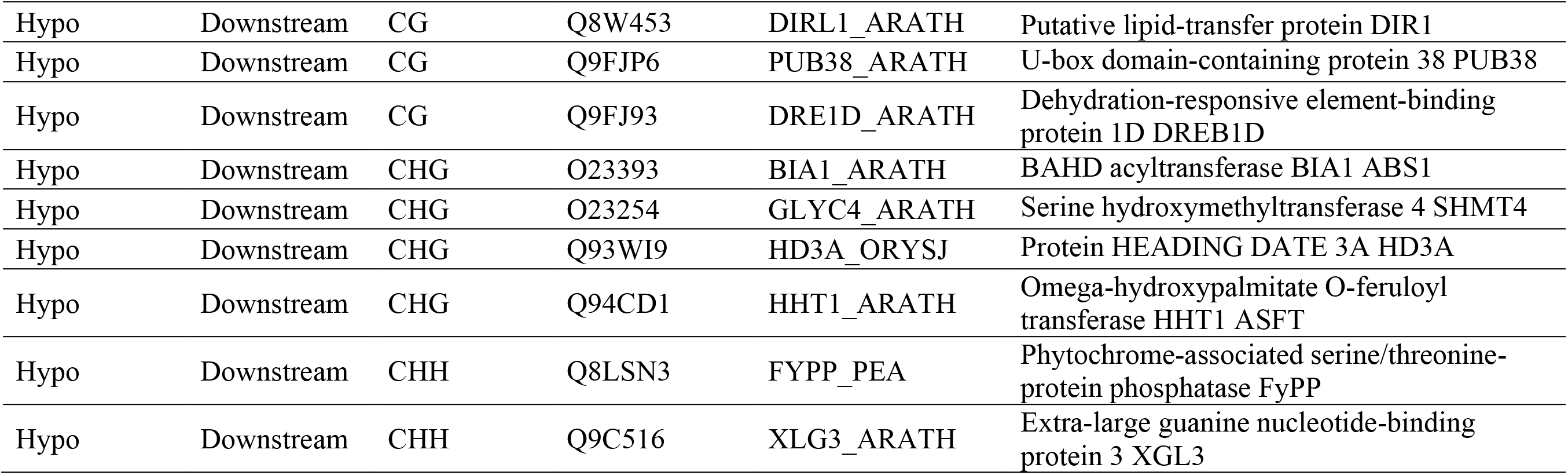
Protein sequences in the UniProtKB Reviewed (Swiss-Prot) database significantly aligning to genes associated with DMRs both hypermethylated and hypomethylated in BF twins compared to No-BF twins (e-value ≤ 0.0001)

A total of 92,188 transcription factor binding sites representing 43 transcription factor families were identified in the hyper- and hypomethylated DMRs associated with the same gene and proximity class using the PlantTFDB (Fig. S8). Significantly enriched transcription factor family binding sites in the DMRs include ERF, Dof, and BBR/BPC (Table 4).

**Table 4.** List of the significantly enriched (adjusted p-value < 0.05) transcription factor family binding sites in the shared DMRs.

| Transcription Factor Family | Binding sites in DMRs | Description |
| --- | --- | --- |
| ERF | 23,245 | Hormone regulation; stress response<sup>1</sup> |
| Dof | 7,629 | Metabolism; phytohormone response; photoperiodic regulation<sup>2</sup> |
| BBR/BPC | 4,014 | Flower development; stem cell size; seed development<sup>3</sup> |
| bHLH | 3,886 | Phytochrome signaling<sup>4</sup> |
| TCP | 3,231 | Branching; flower development; leaf development; cell cycle regulation<sup>5</sup> |
| B3 | 2,518 | Vegetative and reproductive development<sup>6,7</sup> |
| LBD | 2,272 | Leaf, flower, and root development; auxin response<sup>8</sup> |
| GATA | 2,245 | Nitrogen metabolism; stress response; hormonal signaling<sup>9</sup>; meristem development<sup>10</sup> |
| AP2 | 2,061 | Flower development<sup>11</sup> |
| GRAS | 1,209 | Gibberellin signaling; root patterning; light signaling; nodule formation<sup>12</sup> |
| BES1 | 523 | Brassinosteroid regulation<sup>13</sup> |
| E2F/DP | 499 | Cell cycle regulation<sup>14</sup> |
| RAV | 464 | Regulation of flowering; gibberellin regulation<sup>15</sup> |
| C3H | 353 | Plant organ identity; embryogenesis; leaf senescence<sup>16</sup> |
| GeBP | 233 | Leaf primordia development<sup>17</sup> |
| ARR-B | 229 | Cytokinin regulation; shoot development<sup>18</sup> |
| GRF | 80 | Leaf and stem development; flowering; plant longevity<sup>19</sup> |

<sup>1</sup>Xie *et al.*, 2019; <sup>2</sup>Yanagisawa, 2016; <sup>3</sup>Theune *et al.*, 2019; <sup>4</sup>Duek and Fankhauser, 2005; <sup>5</sup>Danisman, 2016; <sup>6</sup>Peng and Weselake, 2013; <sup>7</sup>Swaminathan *et al.*, 2008; <sup>8</sup>Majer and Hochholdinger, 2011; <sup>9</sup>Gupta *et al.*, 2017; <sup>10</sup>Zhao *et al.*, 2004; <sup>11</sup>Okamuro *et al.*, 1997; <sup>12</sup>Hofmann, 2016; <sup>13</sup>Song *et al.*, 2018; <sup>14</sup>Vandepoele *et al.*, 2005; <sup>15</sup>Matías-Hernández *et al.*, 2014; <sup>16</sup>Jiang *et al.*, 2014; <sup>17</sup>Curaba *et al.*, 2003; <sup>18</sup>Xie *et al.*, 2018; <sup>19</sup>Omidbakhshfard *et al.*, 2015

### Transcriptomic analysis of ‘Stukey’ twin pairs

Sequencing was performed to characterize the transcriptome of each ‘Stukey’ twin (Table S5). Of the 25,392 annotated, expressed transcripts detected in the ‘Stukey’ twin pairs (out of the 27,487 annotated transcripts in the ‘Nonpareil’ genome), 16 were found to be significantly differentially expressed with three downregulated in the BF twins and 13 upregulated in the BF twins compared to the no-BF twins (Table 5, Fig. S9). Annotation of these genes revealed involvement in processes such as cell-wall synthesis and metal-ion transport. Principal component analysis of expressed gene counts shows Principal Component (PC) 1 explaining 49% of variance by the separation of twin pairs and PC 2 explaining 45% of variance by the separation of BF condition (Fig. S10).

**Table 5.** Annotation of significantly differentially expressed genes in the BF twins compared to the no-BF twins (p-value < 0.1).

| **Log2**<br>**Fold Change** | **p-adjusted** | **Protein ID** | **Gene Name** |
| --- | --- | --- | --- |
| 2.97 | 4.66E-10 | Undescribed protein |  |
| 2.34 | 0.011 | Oligopeptide transporter 1 | OPT1 (At5g55930) |
| 1.94 | 2.04E-04 | ABC transporter C family member 5 | ABCC5/MRP5 (At1g04120) |
| 1.14 | 0.058 | Putative mediator of RNA<br>polymerase II |  |
| 1.06 | 0.058 | Rust resistance kinase Lr10 | LRK10 |
| 1.05 | 0.058 | Cyclic nucleotide-gated ion channel 1 | CNGC1 (At5g53130) |
| 1.05 | 0.058 | Cellulose synthase-like protein G2 | CSLG2 |
| 0.92 | 0.058 | Early nodulin-like protein 2 | AT4G27520 |
| 0.81 | 0.079 | Cytochrome P450 82A3 | CYP82A3 |
| 0.68 | 0.090 | Protein SAR DEFICIENT 1 | SARD1 (At1g73805) |
| 0.64 | 0.090 | Beta-glucosidase 12 | BGLU12 (OsI_16288) |
| 0.47 | 0.100 | Methionine gamma-lyase | MGL (At1g64660) |
| 0.32 | 0.090 | DMR6-LIKE OXYGENASE 1 | DLO1/SAG108 (At4g10500) |
| -0.59 | 0.090 | Polygalacturonase | PGLR_PRUPE |
| -0.81 | 0.058 | GDSL esterase/lipase | At2g04570 |
| -0.93 | 0.058 | Proline-rich protein 4-like |  |

### Differentially expressed transcripts associated with bud failure exhibition

Expression analysis of DMR-associated genes in each proximity class and methylation context revealed one significantly differentially expressed gene, *CNGC1*, associated with CG hypomethylated intragenic DMRs in both twin pairs. Expression patterns (Log2FoldChange) of DMR-associated genes in both twin pairs do not show a consistent pattern of up- or downregulation separated either by methylation context (Fig. **S5a,b**,c) or proximity class (Fig. **S6a,b,c**).

Interestingly, an uncharacterized protein of 72 amino acids in length showing the highest log2fold change (2.97, Table 5) is located on chromosome 6 in the ‘Nonpareil’ genome. Results using several *in silico* approaches to further analyze this protein suggest it is related to plant development and most likely localized in the nucleus. Three putative motifs sites were identified within the amino acid sequence including one aspartic acid-rich region (19-67) and two Casein kinase II phosphorylation sites (12-15 and 45-48). The protein (C332H463N77O129S3) is composed mostly of aspartic acid (27.8%) and glycine (11.1%) and is predicted to have an overall negative charge with an estimated molecular weight of 7693 kDa. The aliphatic index is predicted to be 59.86 and the average hydropathicity is −0.544. Results from Motif analysis on GenomeNet produced 14 motif alignments, all to entries in the NCBI-CDD database. The most significant alignment (32 amino acids) was to a Cwf15/Cwf15 cell cycle protein from *Schizosaccharomyces pombe* and *S. cerevisiae* (e-value = 0.028, score = 31.4).

## V. Discussion

The goal of this study was to test the hypothesis that DNA methylation signatures are associated with bud failure (BF) exhibition in almond, and to identify and quantify signatures that may have a role in BF-exhibition, emphasizing the almond genome gene space. Previous work representing a first approach to deciphering the methylome landscape in almond showed a lack of independence between BF and DNA methylation; however, genes or genetic regions influenced by DNA methylation that may have a role in BF-exhibition were not elucidated (Fresnedo-Ramírez *et al.*, 2017). The current study represents the first comprehensive scrutiny of the methylome landscape in almond with the purpose of interrogating a nonpathogenic disorder.

### A monozygotic (MZ) twin-based design enables identification of methylomic signatures associated with BF

A MZ twin-based model (Bell and Spector, 2011; Tan *et al.*, 2015) was used in this study to profile DNA methylation in two twin almond pairs discordant for BF-exhibition. To our knowledge, this is the first time this study model had been implemented in a plant species. Using this model, methylation differences were distinguished between each twin pair independently and then compared those differences to find shared divergence between the two twin pairs. The existence of MZ twins in almond and other plant species is documented (Ensign, 1919; Kenworthy *et al.*, 1973; Litz *et al.*, 1982; Martínez-Gómez and Gradziel, 2003; Aleza *et al.*, 2010) and is particularly useful in systems like almond where the production of isogenic or inbred germplasm is complex, such as in obligated outcrossers with systems of self-incompatibility, or when the number of individuals available for sampling or resources are limited (Ushijima *et al.*, 2003). MZ twins are also advantageous over clonally propagated individuals because chronologic versus ontogenetic ages, cofounding variables impacting epigenetic patterns, are not a factor (Gatsuk *et al.*, 1980; Dubrovina and Kiselev, 2016).

### Bud failure exhibition is associated with genome-wide DNA hypomethylation in almond

Patterns discerned from the ‘Stukey’ twin pairs revealed a significant lack of independence between BF-exhibition and DNA methylation, and suggest a putative association between the BF phenotype and non-CG (i.e. CHG and CHH) hypomethylation. This finding supports the hypothesis that epigenetics and chromatin-based mechanisms contribute to the BF disorder and that DNA methylation patterns may inform BF-exhibition (Fresnedo-Ramírez *et al.*, 2017). Bisulfite sequencing enabled quantification of percent methylation in each context (CG, CHG, and CHH) showing that observed methylation falls within levels typically seen in other angiosperms including peach (Niederhuth *et al.*, 2016; Bartels *et al.*, 2018). Observed genome-wide methylation patterns show higher levels of methylation in conserved regions on each chromosome, consistent with patterns in centromeric and pericentromeric regions in other plant species (Fig. S1, Fig. S2, Fig. S3) (Lister *et al.*, 2008; Yan *et al.*, 2010).

There was significant hypomethylation in non-CG contexts in the BF twins, while patterns of hyper- and hypomethylation in the CG context were not as clear. Non-CG methylation is prevalent among plant genomes and can be highly variable, impacting transposition and expression of transposons, early and late-stage plant development, and response to stressors (Kenchanmane Raju *et al.*, 2019). Levels of genome-wide CHH methylation were found to vary seasonally in cotton, resulting in developmental changes and impacting fiber production due to alterations in gene expression (Jin *et al.*, 2013). In the African oil palm, hypomethylation of transposons in the CHG context is associated with the ‘mantling’ phenotype, resulting in desiccated fruit often not observed until the sixth year of growth and causing dramatically reduced oil yields in tissue culture-derived germplasm (Ong-Abdullah *et al.*, 2015). Like mantling in the African oil palm, BF-exhibition in almond can take several years to appear in afflicted trees.

DNA demethylation occurs actively and passively in plants throughout the aging process or after exposure to abiotic and biotic stressors (Liu *et al.*, 2015; Parrilla-Doblas *et al.*, 2019; Liu and Lang, 2020). For example, a study in *Quercus robur* L. demonstrated genome-wide hypomethylation in seeds with advanced age and after exposure to heat stress, resulting in reduced viability (Michalak *et al.*, 2015). BF-exhibition was previously linked to prolonged heat stress in almond (Hellali and Kester, 1979) which could have implications in triggering activation of DNA demethylation mechanisms (Centomani *et al.*, 2015; Liu *et al.*, 2015; Naydenov *et al.*, 2015; Harkess, 2018; Liu *et al.*, 2018). Heat-induced hypomethylation can also result in heritable alterations to DNA methylation profiles including reactivation of transposable elements, as has been demonstrated in Arabidopsis (Lang-Mladek *et al.*, 2010; Ito *et al.*, 2011; Sanchez and Paszkowski, 2014; Liu *et al.*, 2015; Tricker, 2015; Yang *et al.*, 2020), though abiotic stressors do not always induce these heritable marks (Ganguly *et al.*, 2017). Since BF-exhibition can be transmitted by vegetative propagation and sexual reproduction, epigenetic inheritance through chromatin modifications, such as changes in DNA methylation patterns, could serve as a mode of transmission (Kester *et al.*, 1975; Kester *et al.*, 2004). A pedigree study could also be performed to test the inheritance/transmission of differentially methylated regions (DMRs) identified in this study or other chromatin patterns found to be associated with BF-exhibition.

More than 70% of the identified DMRs were classified into one of three proximity classes relative to a gene. Of these DMR-gene associations, permutation testing revealed significantly more DMRs in the up- and downstream proximity classes and significantly fewer in the intragenic proximity class relative to a gene. This result supports previous findings showing gene body methylation in plants tends to be stable over time (and even across multiple generations [Luo *et al.*, 2020]), while methylation occurring up- or downstream of genes tends to exhibit greater dynamism, particularly in response to stressors (Bewick and Schmitz, 2017; Picard and Gehring, 2017; Ito *et al.*, 2019).

### Genes associated with DMRs are involved in meristem development, DNA methylation, dormancy, and response to heat stress

Of the DMR-associated genes, 52 showed significant similarity to previously annotated genes in related species. These genes are significantly enriched in biological processes including protein sumoylation and vegetative meristem development and transition. Identified DMR-associated genes are also involved in meristem maintenance, cell-wall biogenesis, dormancy, and cell-cycle regulation including a probable pectin methylesterase (*PME*), tesmin (*TCX5*), a ribosomal protein (*RPL27)*, an agamous-like MADS-box (*AGL14*), and maintenance of meristems (*MAIN*). These genes and processes represent interesting targets for further functional analysis in almond to explore what, if any, impact they might have on BF development. Available antibodies for the DMR-associated genes could also be used to identify their regulatory functions or to look for active protein expression in key tissues such as meristems.

Empirical evidence suggests a relationship between dormancy processes in the summer (i.e. summer dormancy) and BF-exhibition in almond (Micke, 1996; Kester *et al.*, 2004). Progression of BF results in necrosis of vegetative axillary buds following summer dormancy and occurs by degradation of tunica cells followed by an overgrowth and subsequent collapse of corpus tissues in the meristem (Hellali *et al.*, 1978). Meristems are complex structures with orchestrated growth cycles dependent on internal and external cues (Barton, 2010; Paul *et al.*, 2014; Lloret *et al.*, 2018). Growth and development of meristematic tissues, including dormancy processes, can be regulated by chromatin marks including DNA methylation and histone modifications (Bitonti *et al.*, 2002; Conde *et al.*, 2017; Le Gac *et al.*, 2018; Conde *et al.*, 2019). In poplar, changes in DNA methylation following exposure to heat stress in the previous summer were still present in meristems following winter dormancy, suggesting an epigenetic “memory” (Le Gac *et al.*, 2018). Results herein suggest dysregulation of cell wall biogenesis pathways in almond meristematic tissues could contribute to disruption of dormancy initiation leading to the BF phenotype, and that this alteration could be initiated by abiotic stress (i.e. heat) and maintained in the meristem. DMR-associated genes involved in dormancy or meristem maintenance provide targets for further study to elucidate their involvement in summer dormancy and putative role in the onset of BF in almond (Micke, 1996; Kester *et al.*, 2004; Santamaría *et al.*, 2009; Ríos *et al.*, 2014; Lloret *et al.*, 2018; Fadón *et al.*, 2020).

### Patterns of differential expression identified in genes related to cell wall maintenance and metal ion transport

Callose synthase family gene, *CSLG2* was identified as significantly differentially expressed in this study. *CSL* family members are shown in other plant species to be involved in synthesis of non-cellulosic polysaccharides such as hemicellulose (Richmond and Somerville, 2000; Farrokhi *et al.*, 2006). The DMR-associated gene encoding cyclic nucleotide-gated ion channel 1 (*CNGC1*) was identified as both significantly upregulated in BF twins and as containing an intragenic, shared DMR hypomethylated in BF twins (Table 3; Table 5). Disruption of *CNGC1* in Arabidopsis resulted in a reduction calcium ion accumulation in cells (Ma *et al.*, 2006). Proper regulation of calcium uptake in plants is vital to several developmental processes including meristem organization and cell wall biogenesis (Hepler, 2005; Li *et al.*, 2019). Finally, the putative gene coding for an uncharacterized protein was found to have the highest log2 fold change. Data produced *in silico* surveying various databases and algorithms suggest this putative genomic feature may be shared among other members of the Rosaceae. A highly similar (94.3%) protein of unknown function is reported in the peach gene annotation and the protein identified in this study also shows a high degree of similarity (>70%) with undescribed proteins in *Malus* and *Pyrus*. The three genes represent interesting targets for further study to explore their involvement in BF-exhibition.

### Ethylene responsive factor (ERF) transcription factor family binding sites are enriched in DMRs

Transcription factors (TF) can be sensitive to the presence of DNA methylation at their binding sites, potentially impacting regulation of target genes (Domcke *et al.*, 2015; Héberlé and Bardet, 2019). TF families enriched in DMRs include those with functions related to cell cycle regulation, hormone signaling, plant organ identity, and meristem development (as shown in Table 4). The ERF family represents approximately 25% of the TF binding sites identified. The ERF TF family is involved in stress response and hormone signaling, and members have been shown to regulate growth and development in plants (Licausi *et al.*, 2013; Xie *et al.*, 2019). In pear, another Rosaceous crop, an ERF family TF, which is itself regulated by chromatin marks, was found to be involved in regulating budbreak by activating expression of genes associated with cell division in the apex (Anh Tuan *et al.*, 2016). Results from this work and other studies in apple and poplar support the role of ERF family TFs in dormancy initiation and release in woody perennials (Wisniewski *et al.*, 2015; Busov *et al.*, 2016).

The association between bud failure status and DNA methylation signatures in almond was assessed utilizing a monozygotic twin model with two sets of twin almonds and a whole-genome bisulfite sequencing approach to provide the first comprehensive survey of the almond methylome. The goal was to determine whether the methylome has a role in bud failure exhibition. Results from this work support the hypothesis that genome-wide hypomethylation is associated with bud failure exhibition. The approach utilized in this study allowed identification of several differentially methylated regions between bud failure and bud failure-free twins, providing targets for further investigation. Relevant genomic features identified in this study include genes and transcription factors involved in meristem maintenance, cell cycle control, dormancy, and response to heat stress which might be impacted by chromatin alterations and contribute to the bud failure phenotype. These potential targets can be investigated to evaluate their suitability as biomarkers for bud failure exhibition potential. The availability of biomarkers to screen almond germplasm for bud failure susceptibility would be valuable to breeders, producers, and growers to avoid transmission of the disorder to progeny through breeding and propagation. Elucidation of the mechanisms underlying bud failure development would also improve our understanding of this threatening disorder and provide a basis for addressing and mitigating similar identified or undescribed disorders in other plants.

## VI. Experimental Procedures

### Plant Material

The ‘Stukey’ almond twins are maintained at the Wolfskill Experimental Orchards (Almond Breeding Program, University of California – Davis, Winters, CA) and are comprised of eleven pairs of material half-sib (i.e. the father is unknown) MZ twin trees grown from polyembryonic seed from the cultivar ‘Nonpareil’ (Martínez-Gómez *et al.*, 2002; Martínez-Gómez and Gradziel, 2003). The seeds were planted in 2000, and the trees are monitored annually for BF-exhibition. Two pairs discordant for BF-exhibition before May 2017 were selected for profiling in this study. Leaf samples were collected for DNA extraction in May 2017 from the canopy of each tree and immediately stored in desiccation beads until lyophilization. Samples were shipped on ice to the Ohio Agricultural Research and Development Center (OARDC; The Ohio State University, Wooster, OH, USA) and immediately processed for lyophilization. Following lyophilization, samples were stored at −20 °C until DNA extraction. Leaf material from the same two ‘Stukey’ pairs still showing diverged BF-exhibition was collected in April 2018 for RNA extraction. Leaf samples were harvested from two locations within the canopy of each tree and immediately stored on ice. Samples were shipped on dry ice to the OARDC and stored at −45 °C until sample processing.

### DNA Extraction

High-quality DNA was extracted from leaves following a modified version of the protocol outlined in Lodhi et. al (1994). Briefly, samples were ground with a mortar and pestle in liquid nitrogen, and 100 mg of tissue was added to 1 mL of CTAB buffer (20mM EDTA, 100mM Tris-HCl pH 8.0, 1.4M NaCl, 2.0% CTAB w/v, 2% B-mercaptoethanol v/v). Phase separation was performed using chloroform:isoamyl alcohol (24:1 v/v) and DNA was ethanol precipitated. DNA was RNase (Thermo Scientific™, Waltham, MA) treated according to manufacturer’s instructions, and another phase separation and precipitation was performed. DNA quality and concentration were assessed by gel electrophoresis (1% agarose, 100 Volts, 45 minutes), fluorometry using a Qubit™ 4, and spectrophotometry using a NanoDrop™ 1000 (Thermo Scientific™). Two independent DNA extractions representing technical replicates were performed for each ‘Stukey’ individual.

### RNA Extraction

High-quality RNA was isolated from leaves following a modified version of the protocol outlined in Gambino et. al (2008). Briefly, leaf tissue was ground in liquid nitrogen using mortar and pestle pre-treated with RNaseZap™ Wipes (ThermoFisher Scientific, Waltham, MA). Approximately 150 mg of ground material was added to 900 μL CTAB extraction buffer (2% CTAB, 2.5% PVP-40, 2M NaCl, 100 mM Tris-HCl pH 8.0, 25 mM EDTA pH 8.0, 2% betamercapto-ethanol). Phase separation was performed using chloroform:isoamyl alcohol (24:1), followed by precipitation with 3M lithium chloride. The pellet was resuspended in 500 uL SSTE buffer (10 mM Tris-HCl pH 8.0, 1 mM EDTA pH 8.0, 1% SDS, 1M NaCl) and an additional phase separation was performed. RNA was precipitated with isopropanol and resuspended in 30 uL RNase-free water. RNA was DNase-treated using the DNA-*free*™ DNA Removal Kit (ThermoFisher Scientific) according to the manufacturer’s instructions. RNA quality and concentration were assessed by fluorometry using a Qubit™ 4 RNA HS kit and by electrophoresis using a TapeStation (Agilent, Santa Clara, CA).

### Whole Genome Bisulfite Sequencing Library Preparation and Illumina Sequencing

Whole-genome bisulfite libraries were prepared using the Pico Methyl-Seq Library Prep Kit (Zymo Research, Irvine, CA) according to the manufacturer’s instructions. Eight sequencing libraries were prepared, and each library was uniquely barcoded for multiplexing. The kit provides barcodes for multiplexing six libraries, so two compatible barcodes were synthesized by MilliporeSigma (St. Louis, MO). Library concentration and quality were assessed by fluorometry using a Qubit™ 4 and electrophoresis using a TapeStation (Agilent). Libraries were equimolarly pooled prior to sequencing. Bisulfite libraries were sequenced on one lane of the Illumina^®^ MiSeq platform v. 3 to generate 100 bp single-end reads and re-sequenced on one lane of the Illumina^®^ NextSeq 550 platform to generate 75-bp single-end reads.

### Processing and Alignment of Bisulfite-Sequencing Libraries

Bisulfite sequencing reads were assessed using FastQC v. 0.11.7 (Andrews, 2010) and trimmed using TrimGalore v. 0.6.0 and Cutadapt v. 2.1 with default parameters and an additional 10 base pair 3’ and 5’ prime clip (Krueger, 2016). Reads from the MiSeq and NextSeq runs were then combined into single fasta files for each library. Reads were aligned to the ‘Nonpareil’ almond reference genome, and methylation calls were generated using Bismark v. 0.21.0 (Krueger and Andrews, 2011). Genome coverage, mapping efficiency, and genome-wide methylation levels for each methylation context (CHG, CHH, and CG) were calculated for each ‘Stukey’ individual by combining data from the two technical replicate libraries.

The reference genome used in this study corresponds to the gene space of the almond cultivar ‘Nonpareil’ developed using a combination of Illumina and Hi-C data resulting in eight main scaffolds (N90 = 13.96 Mb)(Fresnedo-Ramírez, 2020). This assembly was functionally annotated using short-read and long-read (Oxford Nanopore, OX4 4DQ, United Kingdom) RNA-seq data and the MAKER pipeline, representing 28,637 gene models. The ‘Nonpareil’ plastid genome was produced by aligning reads to the sequence of the plastid genome NC_034696.1 reported in NCBI.

Plots displaying percent methylation for each twin across each of the eight almond chromosomes were generated using the R package ggplot2 v. 3.3.2 (Wicham 2016). To test for lack of independence between methylation status and bud failure exhibition, a Pearson’s Chi-squared test with Yeat’s continuity correction was conducted using the *chisq.test()* function in R to analyze the number of methylated and unmethylated cytosines detected in each twin pair. An effect size, phi, was calculated for each Pearson’s Chi-squared test using the function *ES.chisq.assoc()* from the package powerAnalysis v. 0.2.1 in R. Finally, conversion efficiencies were calculated for each library by aligning reads to the ‘Nonpareil’ almond chloroplast genome and calculating the percent of methylated cytosines observed. Observed methylation in the chloroplast genome represent non-converted, unmethylated cytosines since all chloroplast cytosines are unmethylated (Fojtová *et al.*, 2001). All analyses were performed using the Ohio Supercomputer Center computing resources (Ohio Supercomputer Center, 1987).

### Identification of Differentially Methylated Regions (DMRs) and Permutation Tests

Differentially methylated loci (DML) were identified for each twin pair and methylation context using the R package DSS (Dispersion Shrinkage for Sequencing data) v. 2.36.0 function *DMLtest()* with smoothing (Wu *et al.*, 2013; Feng *et al.*, 2014; Wu *et al.*, 2015; Park and Wu, 2016). Identified DMLs were then used as input in the function *callDMR()* to identify differentially methylated regions (DMRs) comprised of several DMLs with a p-value threshold set to 0.01 for each twin pair and methylation context. DMR-gene associations were defined by a DMR’s proximity to a nearby gene and classified as either upstream, downstream, or intragenic. Bed files were generated containing the genomic coordinates of almond genes throughout the genome and of all identified DMRs in each twin pair. These files were input to the *window* function in bedtools v. 2.29.2 to identify all DMRs within 2,000 base pairs upstream or downstream of a gene, including intragenic DMRs (Quinlan and Hall, 2010). The observed frequency of DMR-gene associations in each proximity class for each twin pair and methylation context combination were analyzed using a two-tailed permutation test with 1,000 iterations. Bedtools v 2.29.2 *shuffle* function was used to generate a null set of DMR coordinates for each iteration based on observed DMRs and the current almond genome assembly. A histogram of DMR lengths was generated using the R package ggplot2 v. 3.3.2 (Wickham, 2016).

DMRs were considered shared between the two twin pairs if they were associated with the same gene in the same proximity class relative to that gene. The observed frequency of shared DMRs in each methylation context and proximity class were analyzed using a two-tailed permutation test with 1,000 iterations. The null DMR sets generated for each twin pair in the previous permutation were compared to determine frequency of shared DMRs within each set using the definition of shared DMR presented above. Venn diagrams were constructed using the R package VennDiagram v. 1.6.20 displaying the number of identified DMRs in each methylation context in each twin pair and the number of shared DMRs (Chen and Boutros, 2011). A heatmap representing the degree of percent methylation within each shared DMR was generated for each twin in each methylation context and in each proximity class using the R package ComplexHeatmap v. 2.2.0 (Gu *et al.*, 2016).

DMRs were further classified based on the percent methylation difference for each twin pair and only those with the same magnitude of methylation difference (positive or negative) between the BF and no-BF twin in each twin pair were maintained. Shared DMRs with a negative methylation difference were classified as hypomethylated in the BF twins relative to the no-BF twins, and shared DMRs with a positive methylation difference were classified as hypermethylated. Linear regression was performed in R v. 4.0.2 using *lm()* to test for significant correlation between the percent methylation of shared DMRs for the BF twins and for the no-BF twins (R Core Team, 2020). An adjusted R^2^ and p-value were calculated for each regression.

### Annotation of Genes Associated with Shared DMRs and Enrichment Analysis

To obtain a fasta file containing the genomic sequence for genes associated with DMRs in each methylation context and proximity class, samtools v. 1.9 *faidx* was used with both the current almond genome and genomic coordinates for each gene (Li *et al.*, 2009; Li, 2011). Fasta files were queried against a UniProtKB Reviewed (Swiss-Prot) database using BLAST+ v. 2.4.0 with the routine *blastx* and an e-value cutoff of 1×10^−6^ to obtain a list of gene entry identifiers (Camacho *et al.*, 2009; The UniProt Consortium, 2019). Identifiers were uploaded to the Retrieve/ID mapping tool on the UniProt website (https://www.uniprot.org/uploadlists) with options from: UniProtKB AC/ID and to: UniProtKB to obtain gene names and ontology (GO) terms. To perform enrichment analysis of GO terms, a reference map of was generated for genes in the current almond genome using the same method described above. The R package clusterProfiler v. 3.16.1 function *enricher()* was used with the DMR-associated gene list and almond GO term reference map, a p-value cutoff of 0.1, and a Benjamini-Hochberg correction (Yu *et al.*, 2012). Both the list of significantly enriched GO terms identified using *enricher()* and the list of all GO terms present in the DMR-associated gene list were annotated using the *term()* and *termLabel()* functions in the R package rols v. 2.16.4 (Gatto, 2020).

To identify putative transcription factor binding sites within shared DMR sequences, fasta files containing genomic sequence for all shared DMRs were used an input in the Transcription Factor Prediction tool on the Plant Transcription Factor Database (PlantRegMap/PlantTFDB v 5.0; https://planttfdb.cbi.pku.edu.cn/prediction.php) (Jin *et al.*, 2017; Tian *et al.*, 2019). Enrichment analysis was based on the genome-wide frequency of transcription factor binding sites for each family in a reference panel generated using the ‘Nonpareil’ reference genome. To test for significant enrichment, a Fisher’s exact test was performed for each family in R v. 4.0.2 with a Bonferroni correction (R Core Team, 2020).

### mRNA Sequencing Library Preparation and Illumina Sequencing

Ribosomal RNA was depleted from 1 μg of DNase-treated RNA from two technical replicates of each ‘Stukey’ sample using the Invitrogen™ RiboMinus™ Plant Kit for RNA-Seq (ThermoFisher Scientific) according to the manufacturer’s instructions. RNA concentration was determined by fluorometry using a Qubit™ 4 RNA HS kit. To prepare mRNA sequencing libraries, 100 ng of rRNA-depleted RNA from each technical replicate was used as input in the NEBNext^®^ Ultra™ II Directional RNA Library Prep with Sample Purification Beads kit (New England BioLabs^®^ Inc., Ipswich, MA) following instructions provided in Section 5 in the kit manual. Each library was uniquely barcoded using NEBNext^®^ Multiplex Oligos for Illumina^®^ (Index Primers Set 1) (New England BioLabs^®^ Inc.). Library concentration and quality were determined by fluorometry using a Qubit™ 4 and electrophoresis using a TapeStation (Agilent). Libraries were then equimolarly pooled and sequenced on one lane of the Illumina^®^ NextSeq 550 platform to generate 75-bp single-end reads.

### Processing and Alignment of mRNA Sequencing Libraries

mRNA sequencing reads were assessed using FastQC v. 0.11.7 (Andrews, 2010). The reference genome was prepared using STAR v. 2.6.0a with *--runMode genomeGenerate and --genomeSAindexNbases 12* (Dobin *et al.*, 2013). Reads were aligned to the prepared genome using STAR v. 2.6.0a with the following commands: *--alignIntronMin 60 --alignIntronMax 6000 --outFilterScoreMinOverLread 0.3 --outFilterMatchNminOverLread 0.3*. Gene counts were tallied for each library using Subread v. 1.5.0 *featureCounts* with the reference ‘Nonpareil’ gff file (Liao *et al.*, 2013). Processing and alignment pipeline were executed on the Ohio Supercomputer Center computing clusters (Ohio Supercomputer Center, 1987).

### Identifying Differentially Expressed Genes and Integrating Expression Data with DMR-Associated Genes

Differentially expressed genes were identified using a significance cutoff of adjusted p-value < 0.1 in the R package DESeq2 v. 1.26.0 after fitting the model, *~Condition+TwinPair* (Love *et al.*, 2014). Shrunken log_2_ fold changes were generated for the condition contrast using the ‘ashr’ shrinkage estimator (Stephens, 2017). Count data were transformed using the *varianceStabilizingTransformation()* function prior to principal component analysis (Anders and Huber, 2010). Heatmaps were generated using the R package ComplexHeatmap v. 2.2.0 (Gu *et al.*, 2016).

### In silico Analysis of a Hypothetical Protein Identified in mRNA Sequencing

Nucleotide and amino acid sequences for an unidentified protein on chromosome 6 of the ‘Nonpareil’ almond genome with ~3 log_2_fold increase can be found in Data S2. A *blastp* was performed on the amino acid sequence using the UniprotKB reference proteome plus Swiss-Prot with the default parameters (The Uniprot Consortium, 2019). Protein motifs were determined within the sequence using ExPASy ScanProsite (release 2020_02) (Castro *et al.*, 2006; Sigrist *et al.*, 2013). The program SVMProt: Protein Functional Family Prediction was used to predict protein functional families using support vector machine and *k*-nearest neighbors algorithms (Cai *et al.*, 2003). The program ProtParam was used to characterize protein properties including predicted charge, molecular weight, and stability (Gasteiger *et al.*, 2005). To predict subcellular localization, the program YLoc was used (Briesemeister, Rahnenführer, *et al.*, 2010a; Briesemeister, Rahnenüfhrer, *et al.*, 2010b). TargetP-2.0 was used to identify putative peptides (Almagro Armenteros *et al.*, 2019). Finally, the Motif tool on the GenomeNet website (https://www.genome.jp/tools/motif/) was used to search a protein query against several databases to identify putative alignments (Marchler-Bauer *et al.*, 2013; Sigrist *et al.*, 2013; Finn *et al.*, 2014)

## VII. Data Statement and Accession Numbers

The ‘Nonpareil’ almond reference genome fasta file, gff file, and the chloroplast genome fasta file and descriptions of the data can be found at https://www.rosaceae.org/publication_datasets. All sequencing data for this project has been deposited to the NCBI Sequence Read Archive under Bioproject PRJNA668135 including biosamples SAMN16403998, SAMN16403999, SAMN16404000, and SAMN16404001. All code used to perform analyses reported in the manuscript can be found at https://github.com/Fresnedo-Lab/DNA-Methylation-In-Almond-Twins.

## VIII. Acknowledgements

We would like to acknowledge Cheri Nemes for her assistance with wet lab portions of this project. We would also like to acknowledge Andrew Michel and Eric Stockinger for providing edits to later versions of this manuscript. To the Ohio Supercomputer Center for access to computing resources and the Translational Plant Sciences Graduate Program for the fellowship for KMDW. This work was supported by The Ohio State University CFAES-SEEDS program grant # 2019-125, the Almond Board of California Grant HORT35, the U.S. Department of Health and Human Services National Institutes of Health - National Cancer Institute - Cancer Center Support Grant (CCSG) P30CA016058, the USDA National Institute of Food and Agriculture AFRI-EWD Predoctoral Fellowship 2019-67011-29558.

## IX. Short Legends for Supporting Information

**Figure S1.** Genome-wide percent methylation in the CG context across the eight almond chromosomes.

**Figure S2.** Genome-wide percent methylation in the CHG context across the eight almond chromosomes.

**Figure S3.** Genome-wide percent methylation in the CHH context across the eight almond chromosomes.

**Figure S4.** Distribution of length of DMRs in all methylation contexts.

**Figure S5a-c.** Differential cytosine methylation in DMRs and gene expression of associated genes between BF and no-BF ‘Stukey’ twins in each methylation context.

**Figure S6a-c.** Differential cytosine methylation in DMRs and gene expression of associated genes between BF and no-BF ‘Stukey’ twins in each proximity class (upstream, intragenic, and downstream of a gene).

**Figure S7a-f.** Degree of methylation difference at each DMR comparing BF twins and no-BF twins in each proximity class (upstream, intragenic, and downstream of a gene).

**Figure S8.** Number of transcription factor binding sites identified in the shared DMRs in all methylation contexts.

**Figure S9.** Differentially expressed genes comparing BF and no-BF ‘Stukey’ twins.

**Figure S10.** Principal component analysis of differentially expressed genes by twin pair and BF-exhibition.

**Table S1.** Summary of statistics from combination of MiSeq and NextSeq sequencing for each ‘Stukey’ twin.

**Table S2.** Percent cytosine methylation in each methylation context (C = cytosine; G = guanine; H = adenine, thymine, or cytosine) calculated for combined technical replicate libraries for each ‘Stukey’ individual (individuals exhibiting BF are represented in bold).

**Table S3a-c.** Contingency tables displaying association between global DNA methylation in each context and BF-exhibition for each twin pair.

**Table S4a-c.** GO terms associated with significant DMRs.

**Table S5.** Summary statistics from an RNASeq experiment for each ‘Stukey’ twin.

**Data S1.** Genomic coordinates of the DMRs in each twin pair between BF and no-BF ‘Stukey’ twins and associated with the same gene. DMRs are also classified as either hyper- or hypomethylated in each methylation context.

**Data S2.** File containing the nucleotide sequence and amino acid sequence in fasta format of the unidentified protein identified as significantly differentially expressed between BF and no-BF ‘Stukey’ twins.

---

**Table S1.** Combined sequencing results from Illumina MiSeq and NextSeq for ‘Stukey’ libraries. Library refers to the combined technical replicate libraries for each ‘Stukey’ individual. The associated SRA Biosample number is listed to access raw data for this sample on the NCBI SRA repository. Total reads are the number of reads produced in the MiSeq and NextSeq sequencing runs and the depth of coverage represents the coverage based on the size of the ‘Nonpareil’ genome (~250 Mbp). Total aligned reads are the number of reads that aligned to the ‘Nonpareil’ almond genome and mapping efficiency is the percentage of total reads that properly aligned. Coverage aligned represents the depth of coverage for aligned reads based on the size of the ‘Nonpareil’ almond genome (~250 Mbp). Finally, conversion efficiency was calculated based on the conversion rate of the ‘Nonpareil’ chloroplast genome, representing a fully unmethylated sequence.

| **Library** | **SRA Biosample Number** | **Total Reads (Depth)** | **Total Aligned Reads** | **Mapping Efficiency** | **Coverage Aligned** | **Conversion Efficiency** |
| --- | --- | --- | --- | --- | --- | --- |
| ‘Stukey’ 1a | SAMN1640-3998 | 116,099,290 (104X) | 42,474,875 | 36.7% | 38X | 98.8% |
| ‘Stukey’ 1b | SAMN1640-3999 | 123,115,678 (110X) | 47,998,414 | 39.1% | 43X | 98.8% |
| ‘Stukey’ 2a | SAMN1640-4000 | 112,172,115 (100X) | 43,716,467 | 38.9% | 39X | 98.9% |
| ‘Stukey’ 2b | SAMN1640-4001 | 78,347,132 (70X) | 28,476,583 | 36.5% | 25X | 98.7% |

**Table S2.**
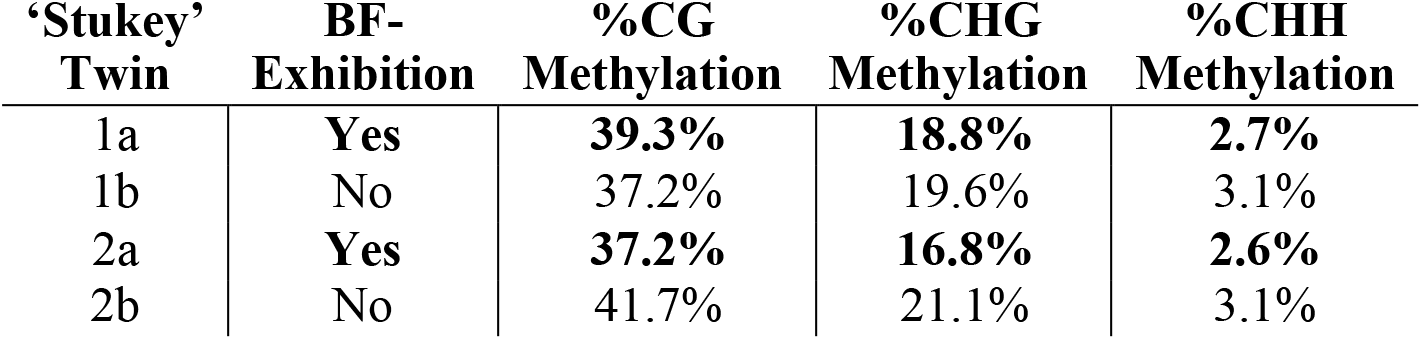
Percent cytosine methylation in each methylation context (C = cytosine; G = guanine; H = adenine, thymine, or cytosine) calculated for combined technical replicate libraries for each ‘Stukey’ individual (individuals exhibiting BF are represented in bold).

| **‘Stukey’**<br>**Twin** | **BF-**<br>**Exhibition** | **%CG**<br>**Methylation** | **%CHG**<br>**Methylation** | **%CHH**<br>**Methylation** |
| --- | --- | --- | --- | --- |
| 1a | **Yes** | **39.3%** | **18.8%** | **2.7%** |
| 1b | No | 37.2% | 19.6% | 3.1% |
| 2a | **Yes** | **37.2%** | **16.8%** | **2.6%** |
| 2b | No | 41.7% | 21.1% | 3.1% |

**Table S3a-c.**
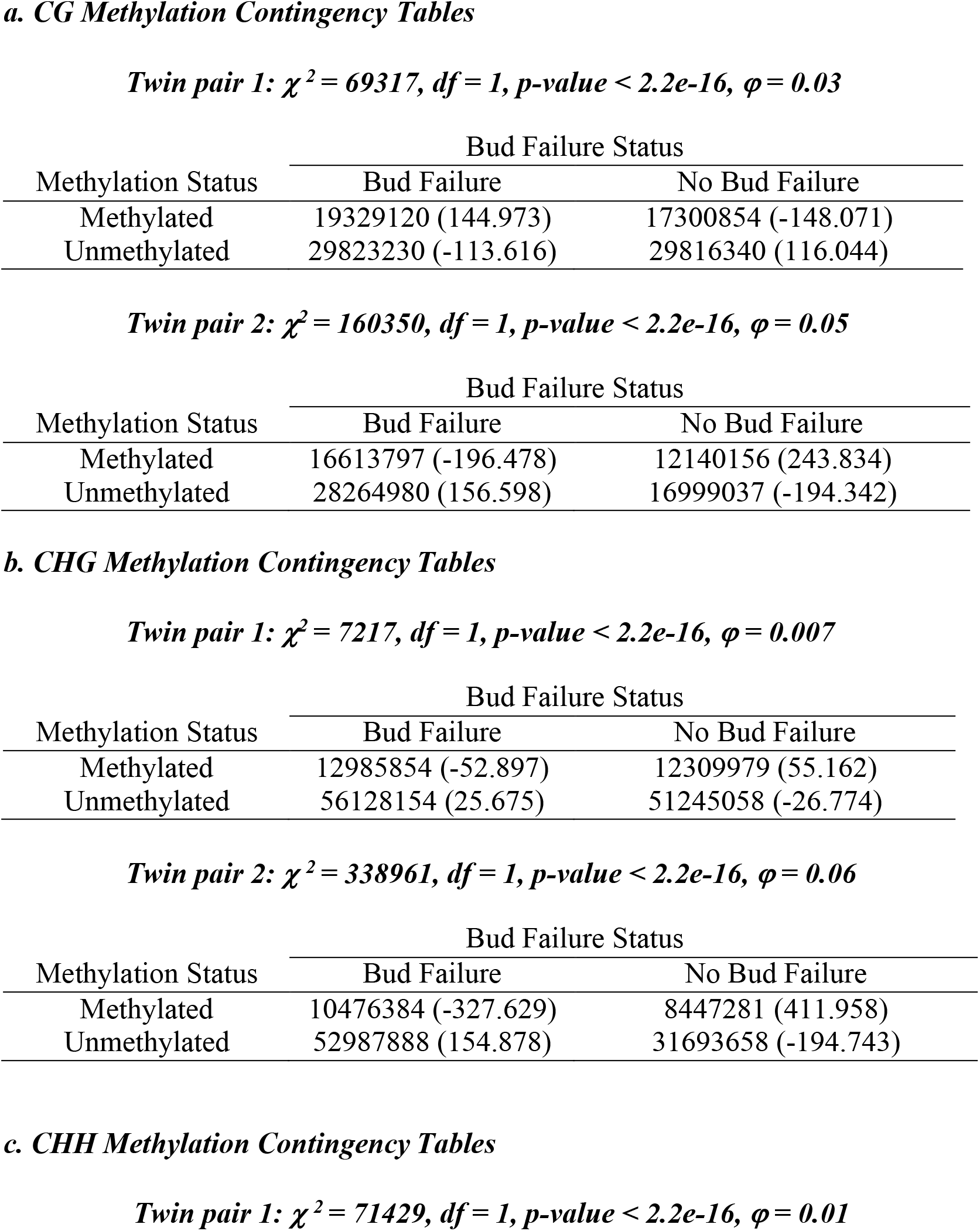
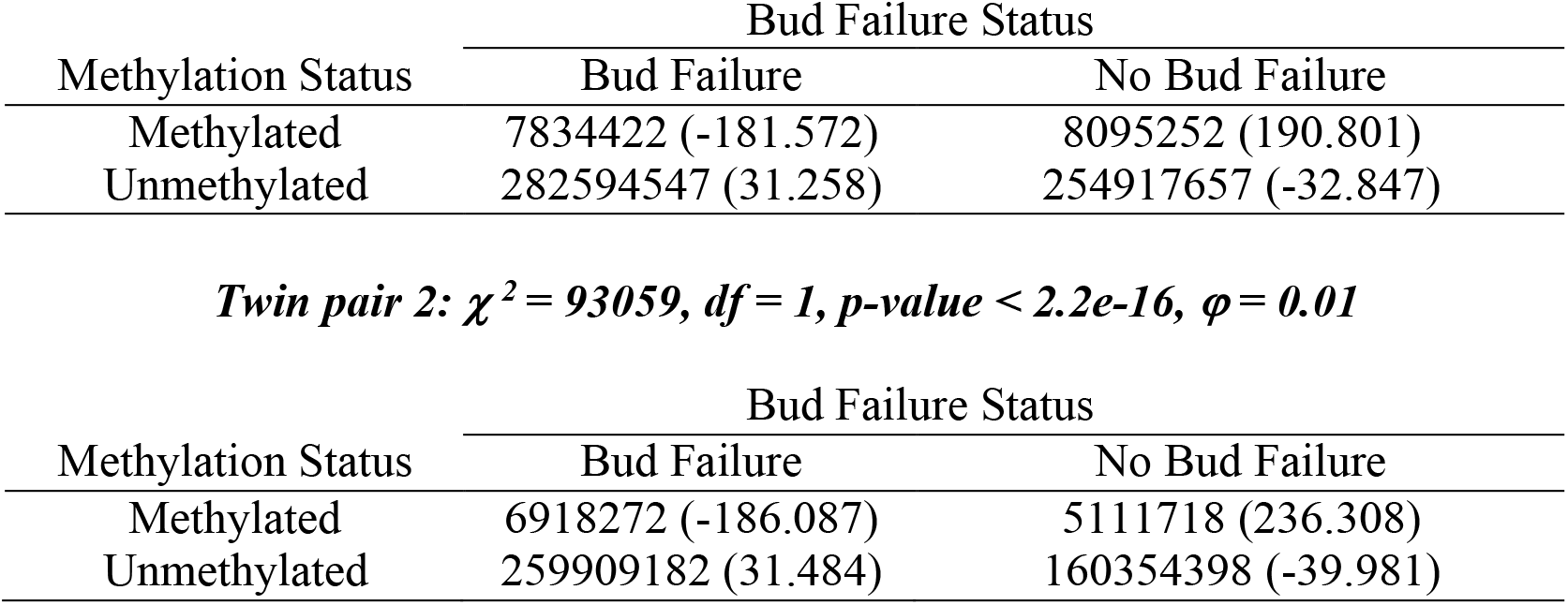
Contingency tables (a), (b), and (c) contain the number of observed methylated and unmethylated cytosines in the CG, CHG, and CHH contexts, respectively, for the BF and no-BF twins in each twin pair (Chi-squared statistic, p-value, and effect size for each comparison is included in the table subtitles and the Pearson’s residual for each count is included in the parentheses in each table).

| Methylation Status | Bud Failure Status |  |
| --- | --- | --- |
|  | Bud Failure | No Bud Failure |
| Methylated | 19329120 (144.973) | 17300854 (-148.071) |
| Unmethylated | 29823230 (-113.616) | 29816340 (116.044) |

***Twin pair 2: $\chi^2 = 160350$ , $df = 1$ , $p\text{-value} < 2.2e-16$ , $\phi = 0.05$***
| Methylation Status | Bud Failure Status |  |
| --- | --- | --- |
|  | Bud Failure | No Bud Failure |
| Methylated | 16613797 (-196.478) | 12140156 (243.834) |
| Unmethylated | 28264980 (156.598) | 16999037 (-194.342) |

**b. CHG Methylation Contingency Tables** ***Twin pair 1: $\chi^2 = 7217$ , $df = 1$ , $p\text{-value} < 2.2e-16$ , $\phi = 0.007$***
| Methylation Status | Bud Failure Status |  |
| --- | --- | --- |
|  | Bud Failure | No Bud Failure |
| Methylated | 12985854 (-52.897) | 12309979 (55.162) |
| Unmethylated | 56128154 (25.675) | 51245058 (-26.774) |

***Twin pair 2: $\chi^2 = 338961$ , $df = 1$ , $p\text{-value} < 2.2e-16$ , $\phi = 0.06$***
| Methylation Status | Bud Failure Status |  |
| --- | --- | --- |
|  | Bud Failure | No Bud Failure |
| Methylated | 10476384 (-327.629) | 8447281 (411.958) |
| Unmethylated | 52987888 (154.878) | 31693658 (-194.743) |

**c. CHH Methylation Contingency Tables** ***Twin pair 1: $\chi^2 = 71429$ , $df = 1$ , $p\text{-value} < 2.2e-16$ , $\phi = 0.01$***
| Methylation Status | Bud Failure Status |  |
| --- | --- | --- |
|  | Bud Failure | No Bud Failure |
| Methylated | 7834422 (-181.572) | 8095252 (190.801) |
| Unmethylated | 282594547 (31.258) | 254917657 (-32.847) |

*Twin pair 2: $\chi^2 = 93059$ , $df = 1$ , $p\text{-value} < 2.2e-16$ , $\phi = 0.01$*
| Methylation Status | Bud Failure Status |  |
| --- | --- | --- |
|  | Bud Failure | No Bud Failure |
| Methylated | 6918272 (-186.087) | 5111718 (236.308) |
| Unmethylated | 259909182 (31.484) | 160354398 (-39.981) |

**Table S4a-c.**
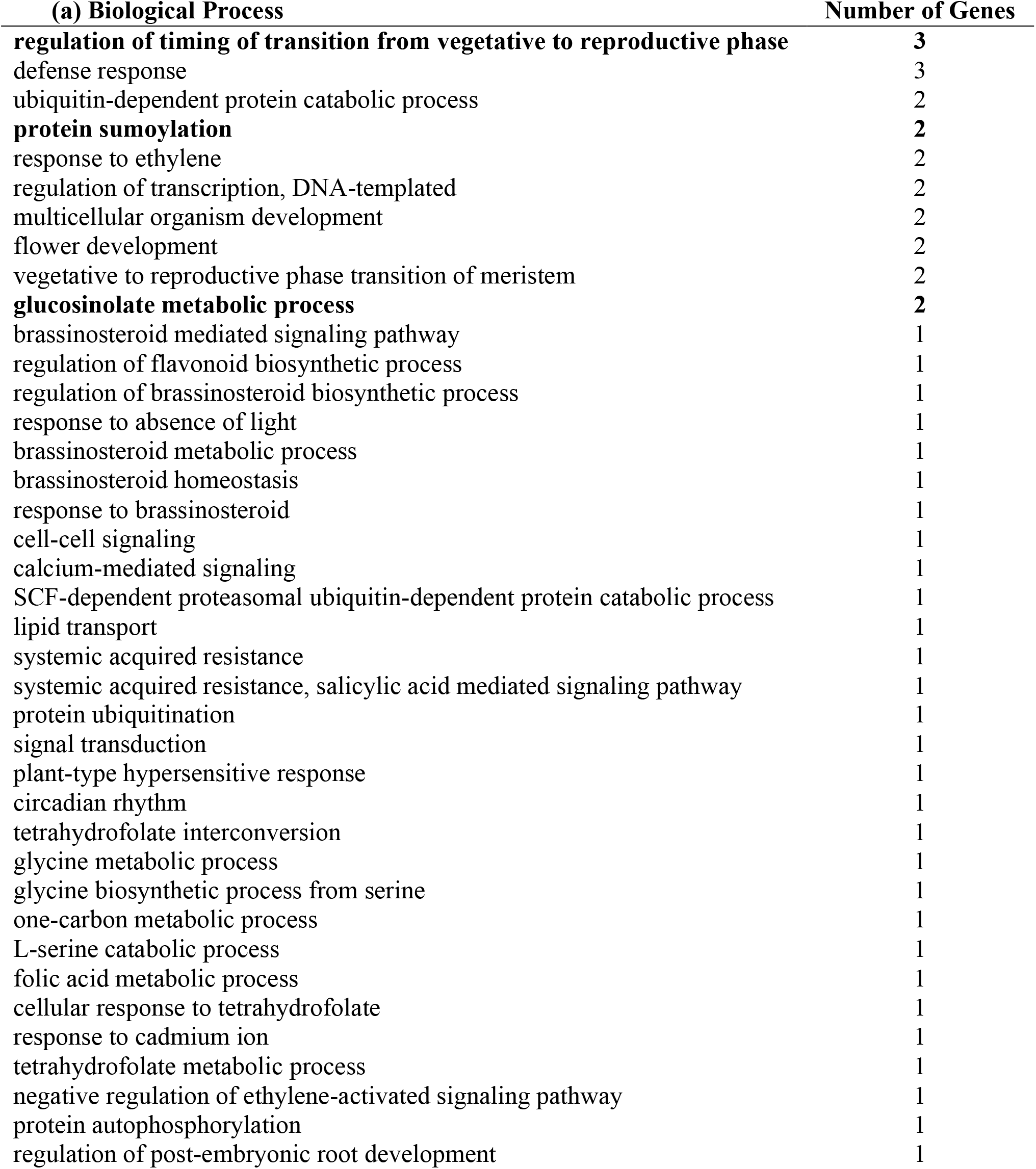
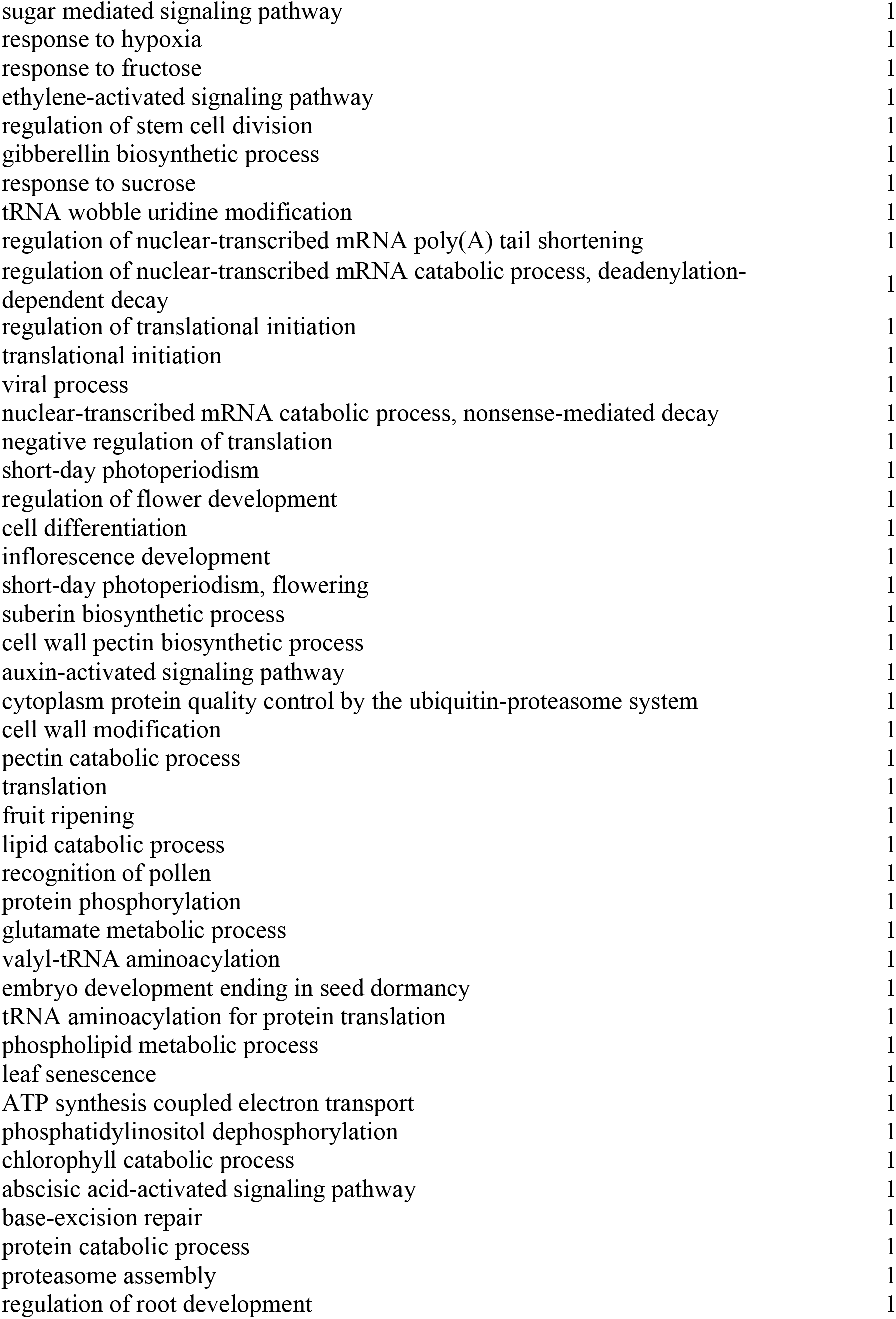
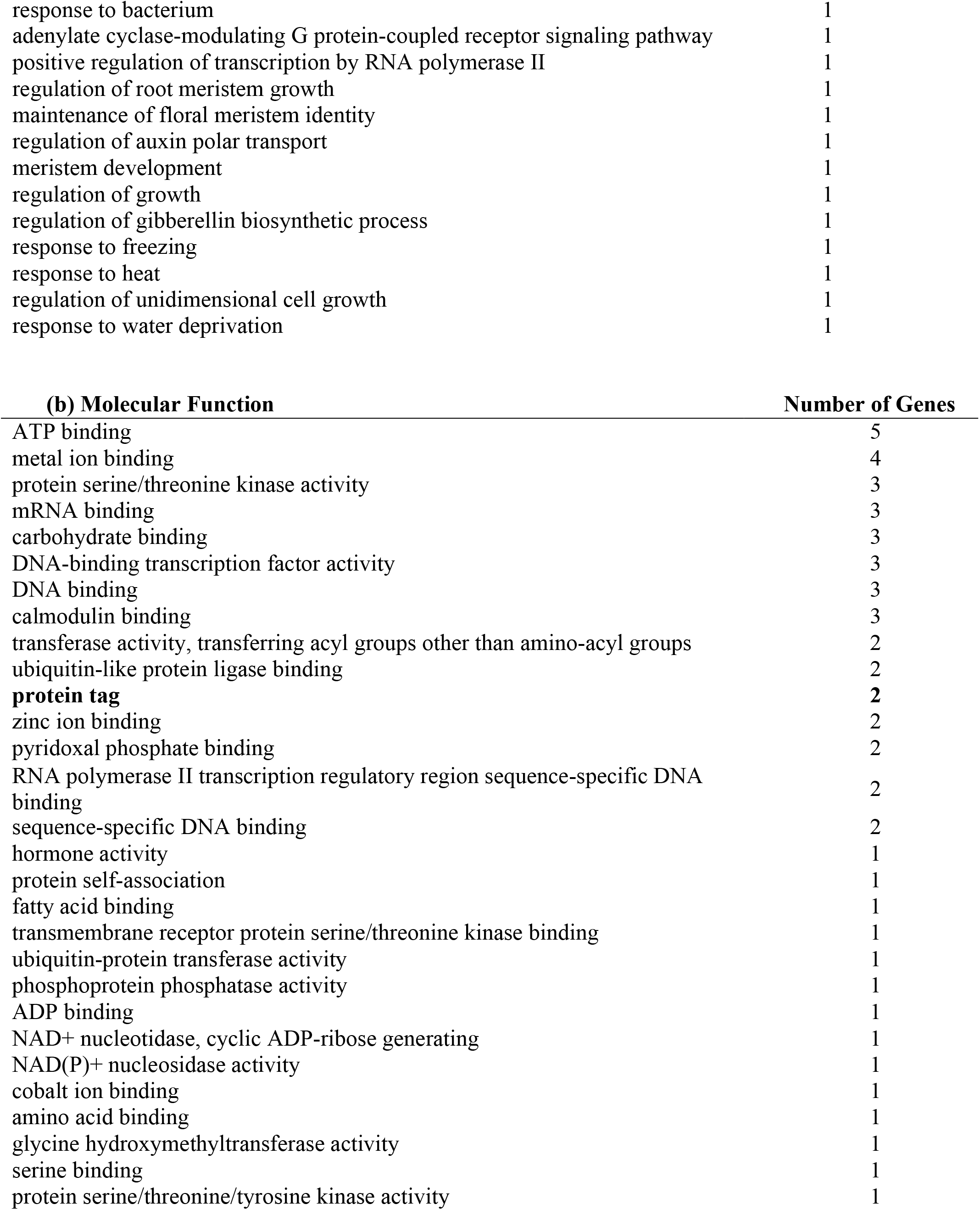
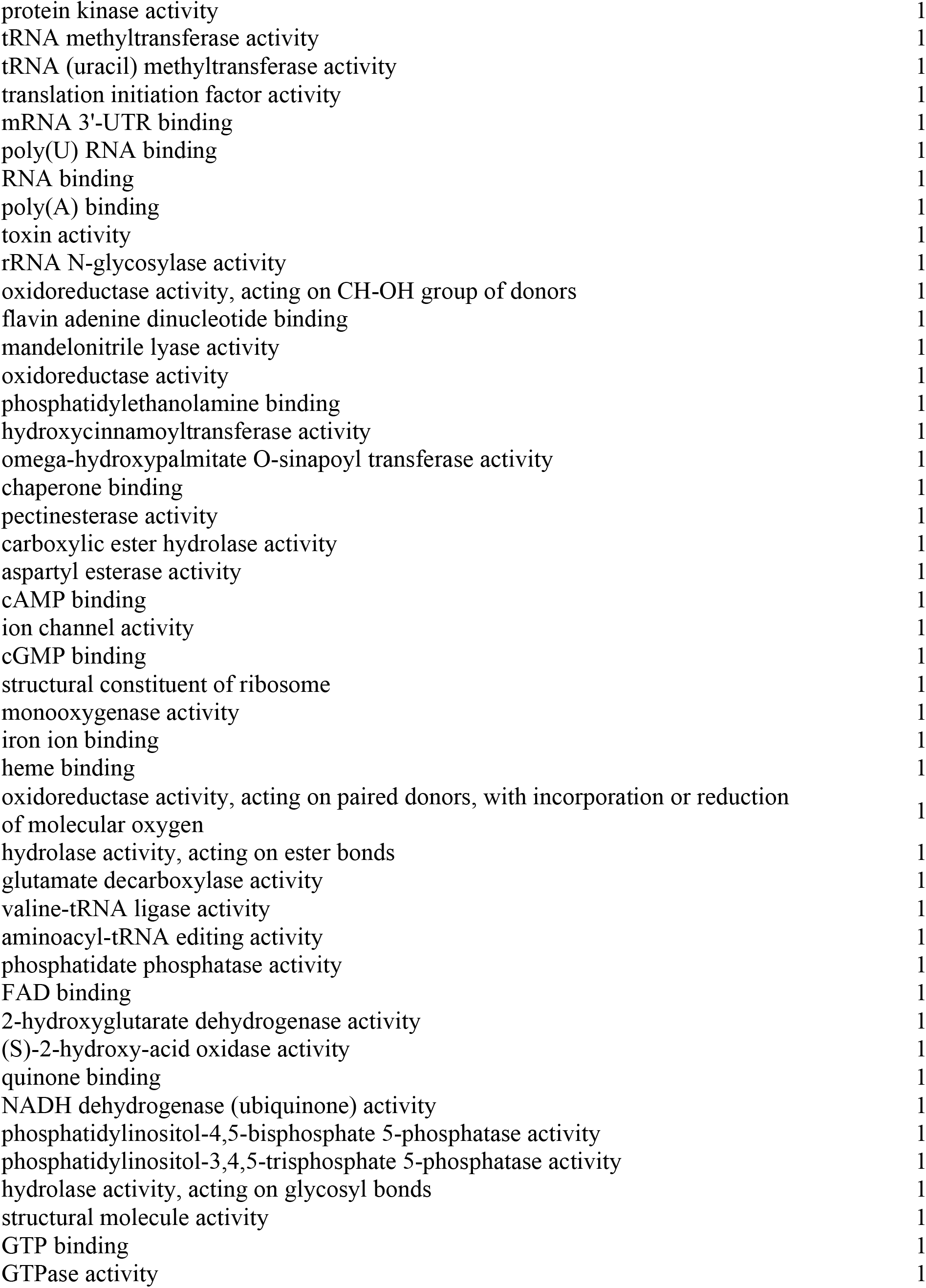
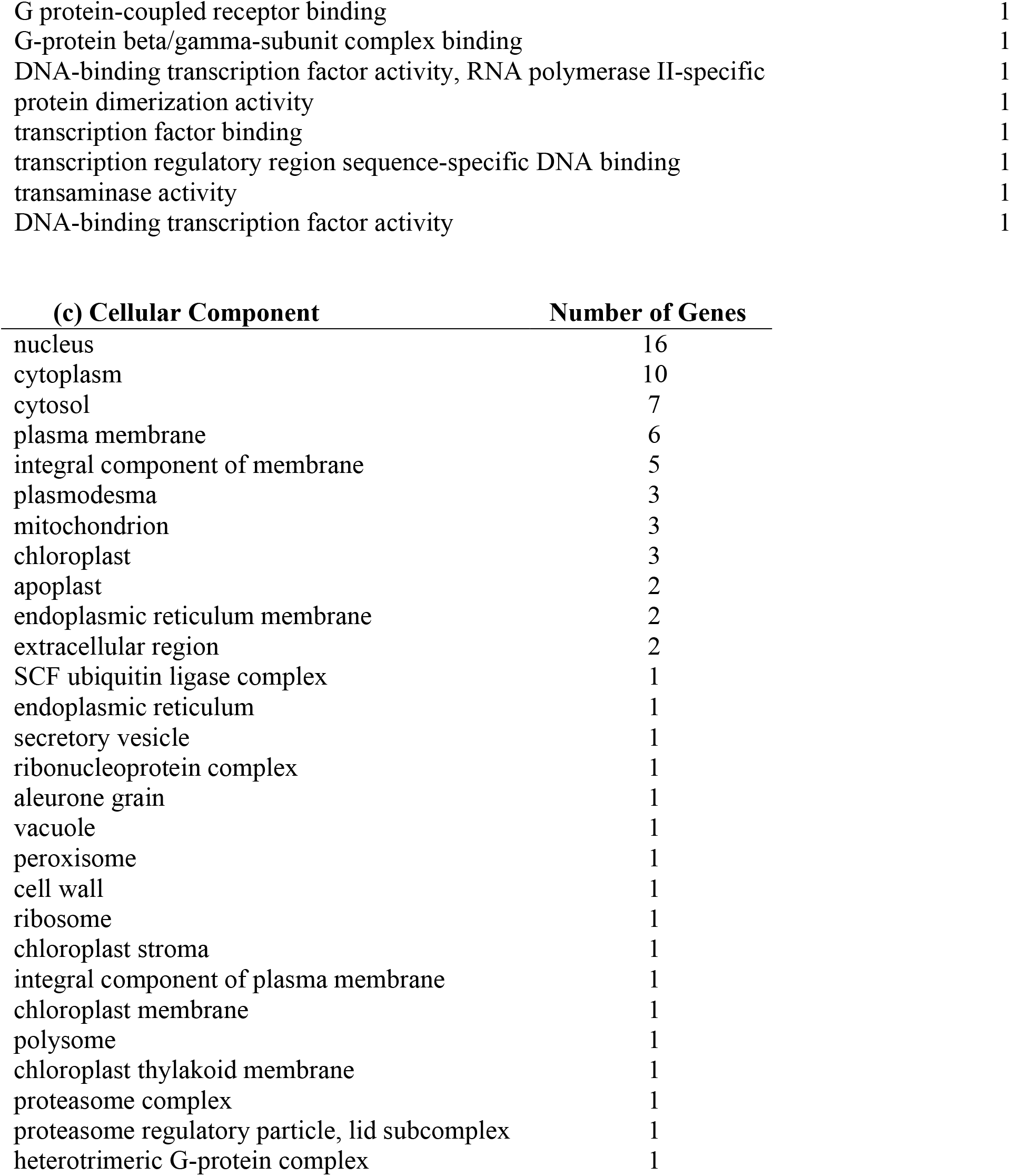
GO terms assigned to DMR-associated genes for each GO category: biological process (a), molecular function (b), and cellular component (c). Significantly enriched GO terms are represented in bold (alpha = 0.1).

| **(a) Biological Process** | **Number of Genes** |
| --- | --- |
| **regulation of timing of transition from vegetative to reproductive phase** | **3** |
| defense response | 3 |
| ubiquitin-dependent protein catabolic process | 2 |
| **protein sumoylation** | **2** |
| response to ethylene | 2 |
| regulation of transcription, DNA-templated | 2 |
| multicellular organism development | 2 |
| flower development | 2 |
| vegetative to reproductive phase transition of meristem | 2 |
| **glucosinolate metabolic process** | **2** |
| brassinosteroid mediated signaling pathway | 1 |
| regulation of flavonoid biosynthetic process | 1 |
| regulation of brassinosteroid biosynthetic process | 1 |
| response to absence of light | 1 |
| brassinosteroid metabolic process | 1 |
| brassinosteroid homeostasis | 1 |
| response to brassinosteroid | 1 |
| cell-cell signaling | 1 |
| calcium-mediated signaling | 1 |
| SCF-dependent proteasomal ubiquitin-dependent protein catabolic process | 1 |
| lipid transport | 1 |
| systemic acquired resistance | 1 |
| systemic acquired resistance, salicylic acid mediated signaling pathway | 1 |
| protein ubiquitination | 1 |
| signal transduction | 1 |
| plant-type hypersensitive response | 1 |
| circadian rhythm | 1 |
| tetrahydrofolate interconversion | 1 |
| glycine metabolic process | 1 |
| glycine biosynthetic process from serine | 1 |
| one-carbon metabolic process | 1 |
| L-serine catabolic process | 1 |
| folic acid metabolic process | 1 |
| cellular response to tetrahydrofolate | 1 |
| response to cadmium ion | 1 |
| tetrahydrofolate metabolic process | 1 |
| negative regulation of ethylene-activated signaling pathway | 1 |
| protein autophosphorylation | 1 |
| regulation of post-embryonic root development | 1 |
| sugar mediated signaling pathway | 1 |
| response to hypoxia | 1 |
| response to fructose | 1 |
| ethylene-activated signaling pathway | 1 |
| regulation of stem cell division | 1 |
| gibberellin biosynthetic process | 1 |
| response to sucrose | 1 |
| tRNA wobble uridine modification | 1 |
| regulation of nuclear-transcribed mRNA poly(A) tail shortening | 1 |
| regulation of nuclear-transcribed mRNA catabolic process, deadenylation-dependent decay | 1 |
| regulation of translational initiation | 1 |
| translational initiation | 1 |
| viral process | 1 |
| nuclear-transcribed mRNA catabolic process, nonsense-mediated decay | 1 |
| negative regulation of translation | 1 |
| short-day photoperiodism | 1 |
| regulation of flower development | 1 |
| cell differentiation | 1 |
| inflorescence development | 1 |
| short-day photoperiodism, flowering | 1 |
| suberin biosynthetic process | 1 |
| cell wall pectin biosynthetic process | 1 |
| auxin-activated signaling pathway | 1 |
| cytoplasm protein quality control by the ubiquitin-proteasome system | 1 |
| cell wall modification | 1 |
| pectin catabolic process | 1 |
| translation | 1 |
| fruit ripening | 1 |
| lipid catabolic process | 1 |
| recognition of pollen | 1 |
| protein phosphorylation | 1 |
| glutamate metabolic process | 1 |
| valyl-tRNA aminoacylation | 1 |
| embryo development ending in seed dormancy | 1 |
| tRNA aminoacylation for protein translation | 1 |
| phospholipid metabolic process | 1 |
| leaf senescence | 1 |
| ATP synthesis coupled electron transport | 1 |
| phosphatidylinositol dephosphorylation | 1 |
| chlorophyll catabolic process | 1 |
| abscisic acid-activated signaling pathway | 1 |
| base-excision repair | 1 |
| protein catabolic process | 1 |
| proteasome assembly | 1 |
| regulation of root development | 1 |
| response to bacterium | 1 |
| adenylate cyclase-modulating G protein-coupled receptor signaling pathway | 1 |
| positive regulation of transcription by RNA polymerase II | 1 |
| regulation of root meristem growth | 1 |
| maintenance of floral meristem identity | 1 |
| regulation of auxin polar transport | 1 |
| meristem development | 1 |
| regulation of growth | 1 |
| regulation of gibberellin biosynthetic process | 1 |
| response to freezing | 1 |
| response to heat | 1 |
| regulation of unidimensional cell growth | 1 |
| response to water deprivation | 1 |

| **(b) Molecular Function** | **Number of Genes** |
| --- | --- |
| ATP binding | 5 |
| metal ion binding | 4 |
| protein serine/threonine kinase activity | 3 |
| mRNA binding | 3 |
| carbohydrate binding | 3 |
| DNA-binding transcription factor activity | 3 |
| DNA binding | 3 |
| calmodulin binding | 3 |
| transferase activity, transferring acyl groups other than amino-acyl groups | 2 |
| ubiquitin-like protein ligase binding | 2 |
| **protein tag** | **2** |
| zinc ion binding | 2 |
| pyridoxal phosphate binding | 2 |
| RNA polymerase II transcription regulatory region sequence-specific DNA binding | 2 |
| sequence-specific DNA binding | 2 |
| hormone activity | 1 |
| protein self-association | 1 |
| fatty acid binding | 1 |
| transmembrane receptor protein serine/threonine kinase binding | 1 |
| ubiquitin-protein transferase activity | 1 |
| phosphoprotein phosphatase activity | 1 |
| ADP binding | 1 |
| NAD<sup>+</sup> nucleotidase, cyclic ADP-ribose generating | 1 |
| NAD(P)<sup>+</sup> nucleosidase activity | 1 |
| cobalt ion binding | 1 |
| amino acid binding | 1 |
| glycine hydroxymethyltransferase activity | 1 |
| serine binding | 1 |
| protein serine/threonine/tyrosine kinase activity | 1 |
| protein kinase activity | 1 |
| tRNA methyltransferase activity | 1 |
| tRNA (uracil) methyltransferase activity | 1 |
| translation initiation factor activity | 1 |
| mRNA 3'-UTR binding | 1 |
| poly(U) RNA binding | 1 |
| RNA binding | 1 |
| poly(A) binding | 1 |
| toxin activity | 1 |
| rRNA N-glycosylase activity | 1 |
| oxidoreductase activity, acting on CH-OH group of donors | 1 |
| flavin adenine dinucleotide binding | 1 |
| mandelonitrile lyase activity | 1 |
| oxidoreductase activity | 1 |
| phosphatidylethanolamine binding | 1 |
| hydroxycinnamoyltransferase activity | 1 |
| omega-hydroxypalmitate O-sinapoyl transferase activity | 1 |
| chaperone binding | 1 |
| pectinesterase activity | 1 |
| carboxylic ester hydrolase activity | 1 |
| aspartyl esterase activity | 1 |
| cAMP binding | 1 |
| ion channel activity | 1 |
| cGMP binding | 1 |
| structural constituent of ribosome | 1 |
| monooxygenase activity | 1 |
| iron ion binding | 1 |
| heme binding | 1 |
| oxidoreductase activity, acting on paired donors, with incorporation or reduction of molecular oxygen | 1 |
| hydrolase activity, acting on ester bonds | 1 |
| glutamate decarboxylase activity | 1 |
| valine-tRNA ligase activity | 1 |
| aminoacyl-tRNA editing activity | 1 |
| phosphatidate phosphatase activity | 1 |
| FAD binding | 1 |
| 2-hydroxyglutarate dehydrogenase activity | 1 |
| (S)-2-hydroxy-acid oxidase activity | 1 |
| quinone binding | 1 |
| NADH dehydrogenase (ubiquinone) activity | 1 |
| phosphatidylinositol-4,5-bisphosphate 5-phosphatase activity | 1 |
| phosphatidylinositol-3,4,5-trisphosphate 5-phosphatase activity | 1 |
| hydrolase activity, acting on glycosyl bonds | 1 |
| structural molecule activity | 1 |
| GTP binding | 1 |
| GTPase activity | 1 |
| G protein-coupled receptor binding | 1 |
| G-protein beta/gamma-subunit complex binding | 1 |
| DNA-binding transcription factor activity, RNA polymerase II-specific | 1 |
| protein dimerization activity | 1 |
| transcription factor binding | 1 |
| transcription regulatory region sequence-specific DNA binding | 1 |
| transaminase activity | 1 |
| DNA-binding transcription factor activity | 1 |

| **(c) Cellular Component** | **Number of Genes** |
| --- | --- |
| nucleus | 16 |
| cytoplasm | 10 |
| cytosol | 7 |
| plasma membrane | 6 |
| integral component of membrane | 5 |
| plasmodesma | 3 |
| mitochondrion | 3 |
| chloroplast | 3 |
| apoplast | 2 |
| endoplasmic reticulum membrane | 2 |
| extracellular region | 2 |
| SCF ubiquitin ligase complex | 1 |
| endoplasmic reticulum | 1 |
| secretory vesicle | 1 |
| ribonucleoprotein complex | 1 |
| aleurone grain | 1 |
| vacuole | 1 |
| peroxisome | 1 |
| cell wall | 1 |
| ribosome | 1 |
| chloroplast stroma | 1 |
| integral component of plasma membrane | 1 |
| chloroplast membrane | 1 |
| polysome | 1 |
| chloroplast thylakoid membrane | 1 |
| proteasome complex | 1 |
| proteasome regulatory particle, lid subcomplex | 1 |
| heterotrimeric G-protein complex | 1 |

**Table S5.** Sequencing results from an RNASeq experiment performed on two ‘Stukey’ twin pairs displaying divergent bud failure exhibition.

| **Library** | **SRA Biosample Number** | **Total Reads Generated** | **Total Reads Aligned** | **Mapping Efficiency** |
| --- | --- | --- | --- | --- |
| ‘Stukey’ 1a | SAMN1640-3998 | 84,640,142 | 56,935,045 | 67.3% |
| ‘Stukey’ 1b | SAMN1640-3999 | 103,419,251 | 69,651,011 | 67.3% |
| ‘Stukey’ 2a | SAMN1640-4000 | 99,578,512 | 66,298,129 | 66.6% |
| ‘Stukey’ 2b | SAMN1640-4001 | 85,576,260 | 60,312,161 | 70.5% |

**Fig. S1.**
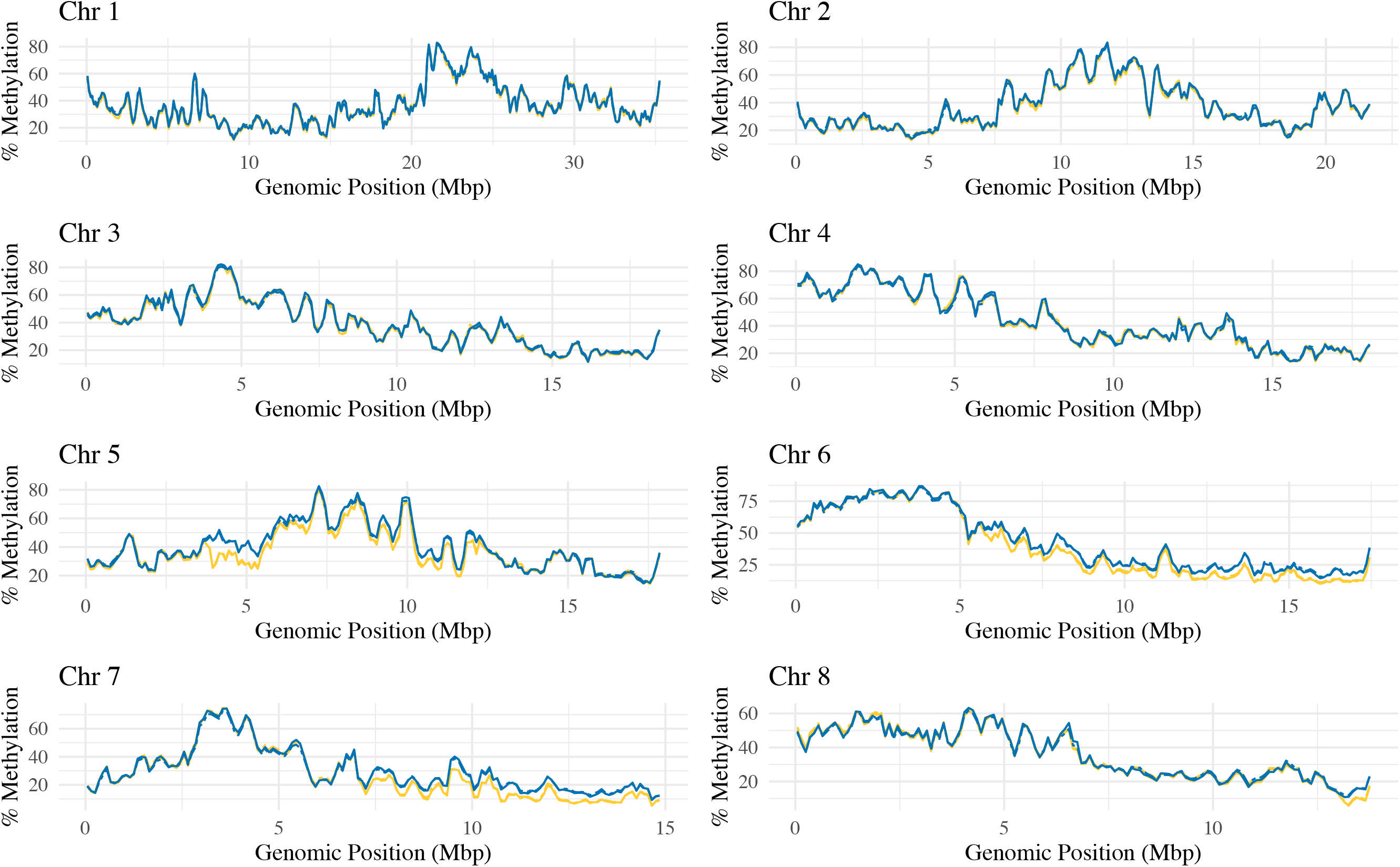
Percent methylation in the CG context across each of the eight ‘Nonpareil’ genome scaffolds representing the eight almond chromosomes. Chromosome number is listed above each plot. The solid lines represent BF individuals and the dashed lines represent no-BF individuals. The gold lines represent ‘Stukey’ twin pair 1 and the blue lines represent ‘Stukey’ twin pair 2.

**Fig. S2.**
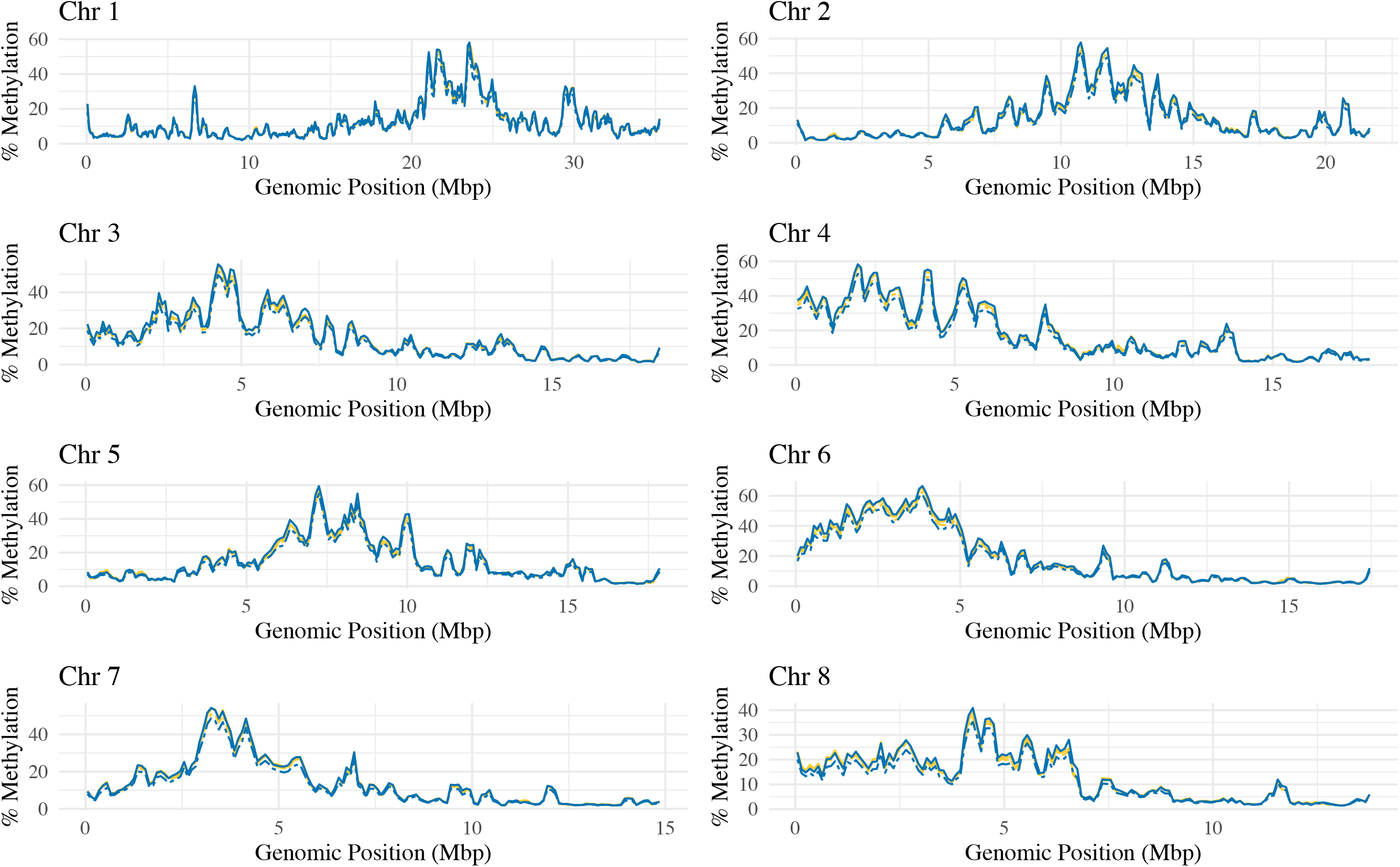
Percent methylation in the CHG context across each of the eight ‘Nonpareil’ genome scaffolds representing the eight almond chromosomes. Chromosome number is listed above each plot. The solid lines represent BF individuals and the dashed lines represent no-BF individuals. The gold lines represent ‘Stukey’ twin pair 1 and the blue lines represent ‘Stukey’ twin pair 2.

**Fig. S3.**
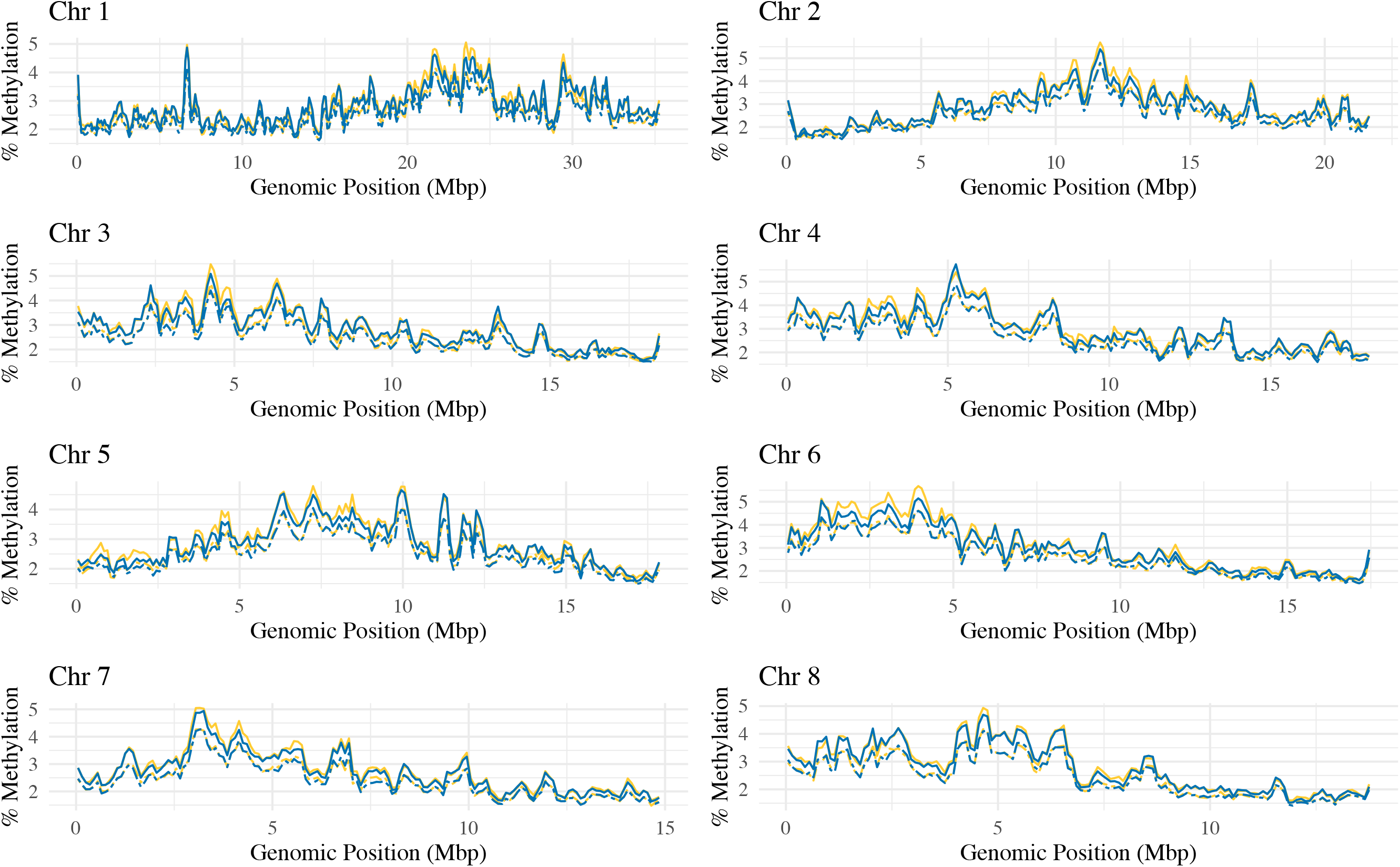
Percent methylation in the CHH context across each of the eight ‘Nonpareil’ genome scaffolds representing the eight almond chromosomes. Chromosome number is listed above each plot. The solid lines represent BF individuals and the dashed lines represent no-BF individuals. The gold lines represent ‘Stukey’ twin pair 1 and the blue lines represent ‘Stukey’ twin pair 2.

**Fig. S4.**
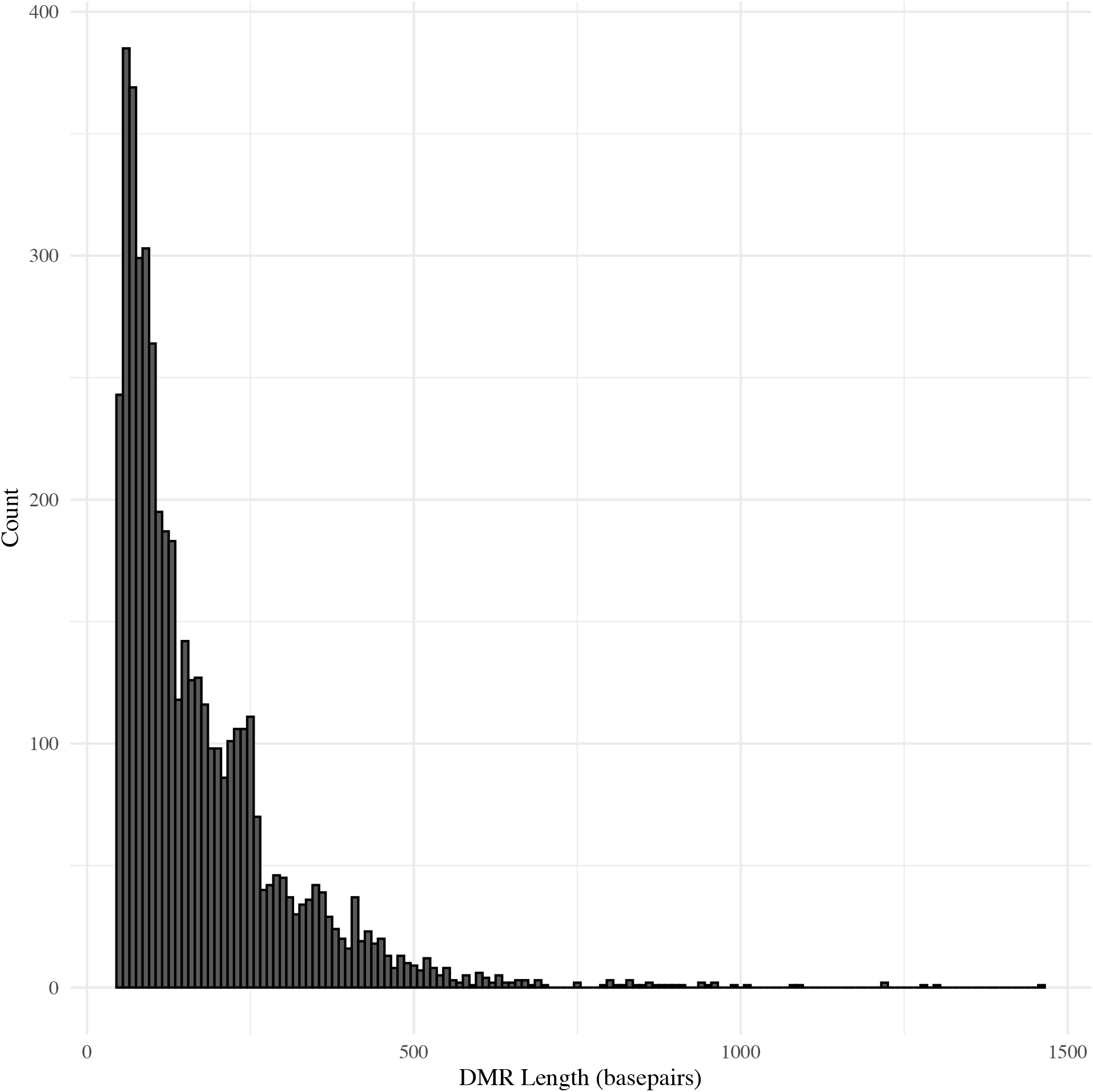
Distribution of length of all DMRs found in each twin pair in all methylation contexts.

**Fig. S5a-c.**
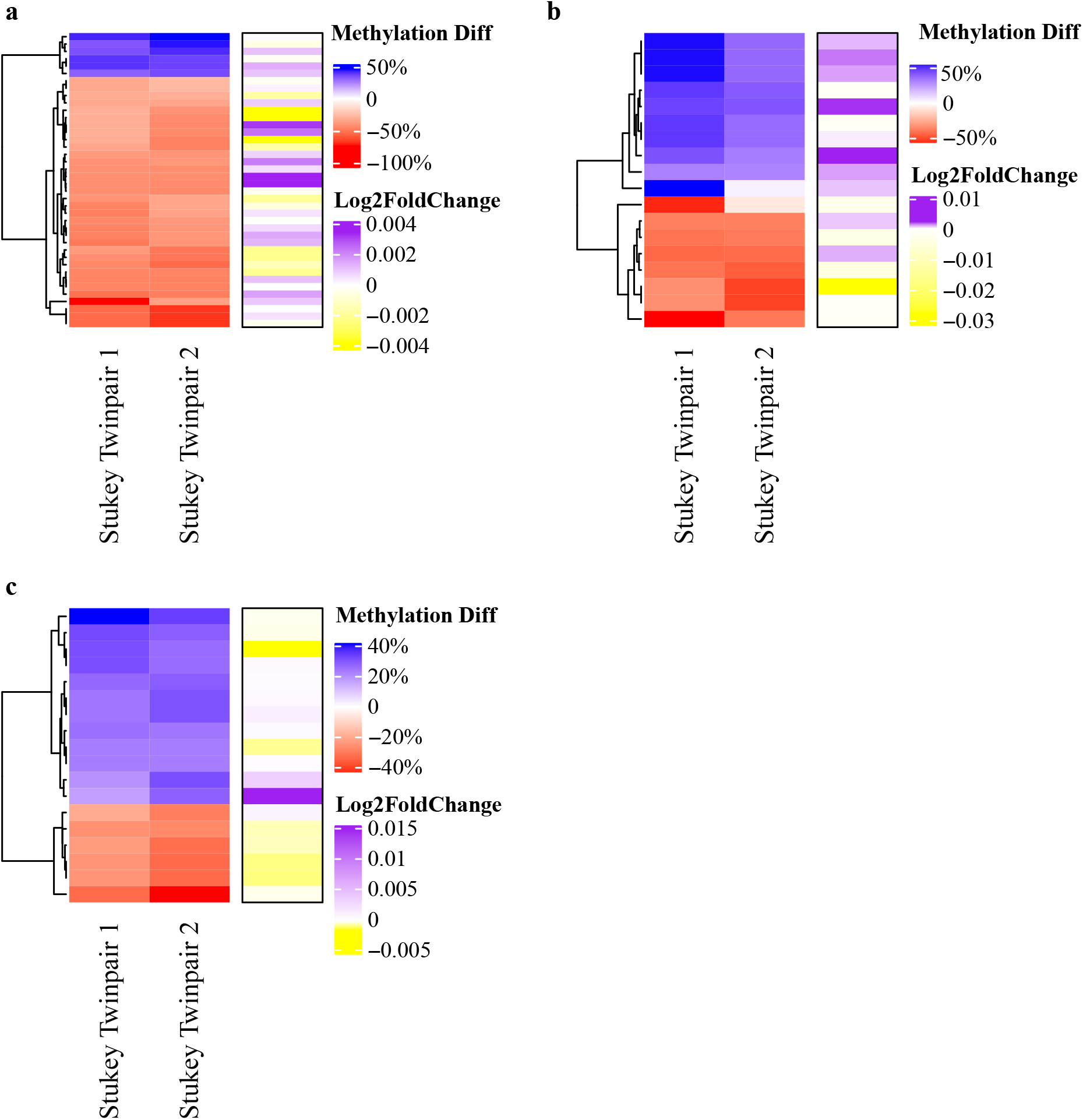
Differential cytosine methylation in each DMR and expression of genes associated with the DMR in almond twin pairs discordant for BF-exhibition. The heatmap in red and blue represents the difference in percent methylation in the BF twin compared to the no-BF twin for every significant shared DMR in each context. The heatmaps in purple and yellow represent the differential expression in the BF twins compared to the no-BF twins for the genes associated with each DMR. Panel **(a)** represents the CG context DMRs, panel **(b)** represents CHG context DMRs, and panel **(c)** represents CHH context DMRs.

**Fig. S6a-c.**
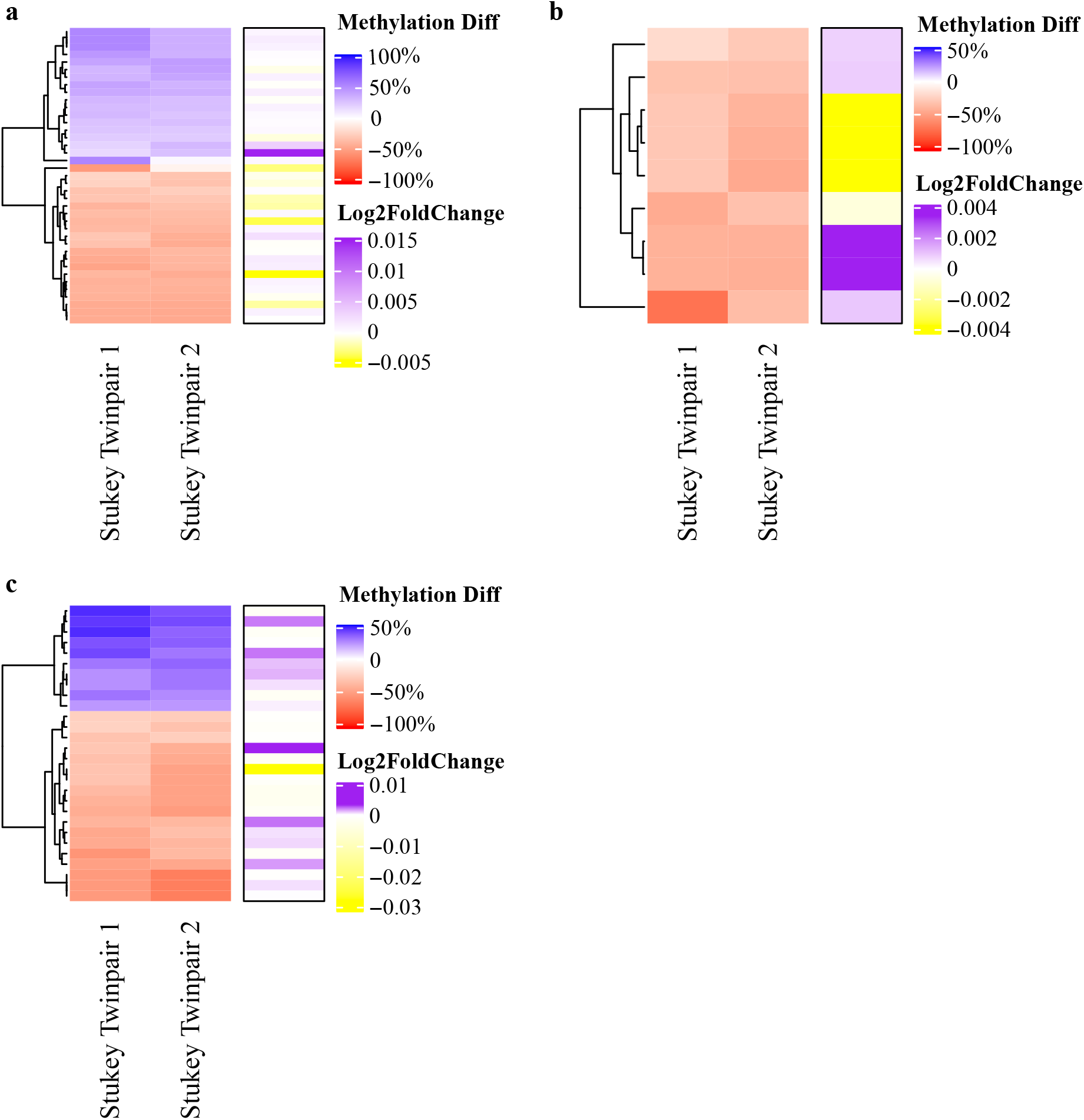
Differential cytosine methylation in each DMR and expression of genes associated with the DMR in almond twin pairs discordant for BF-exhibition. The heatmap in red and blue represents the difference in percent methylation in the BF twin compared to the no-BF twin for every significant shared DMR in each proximity class. The heatmaps in purple and yellow represent the differential expression in the BF twins compared to the no-BF twins for the genes associated with each DMR. Panel **(a)** represents the DMRs upstream (within 2,000 bp) of a gene, panel **(b)** represents the intragenic DMRs, and panel **(c)** represents the DMRs downstream (within 2,000 bp) of a gene.

**Fig. S7a-f.**
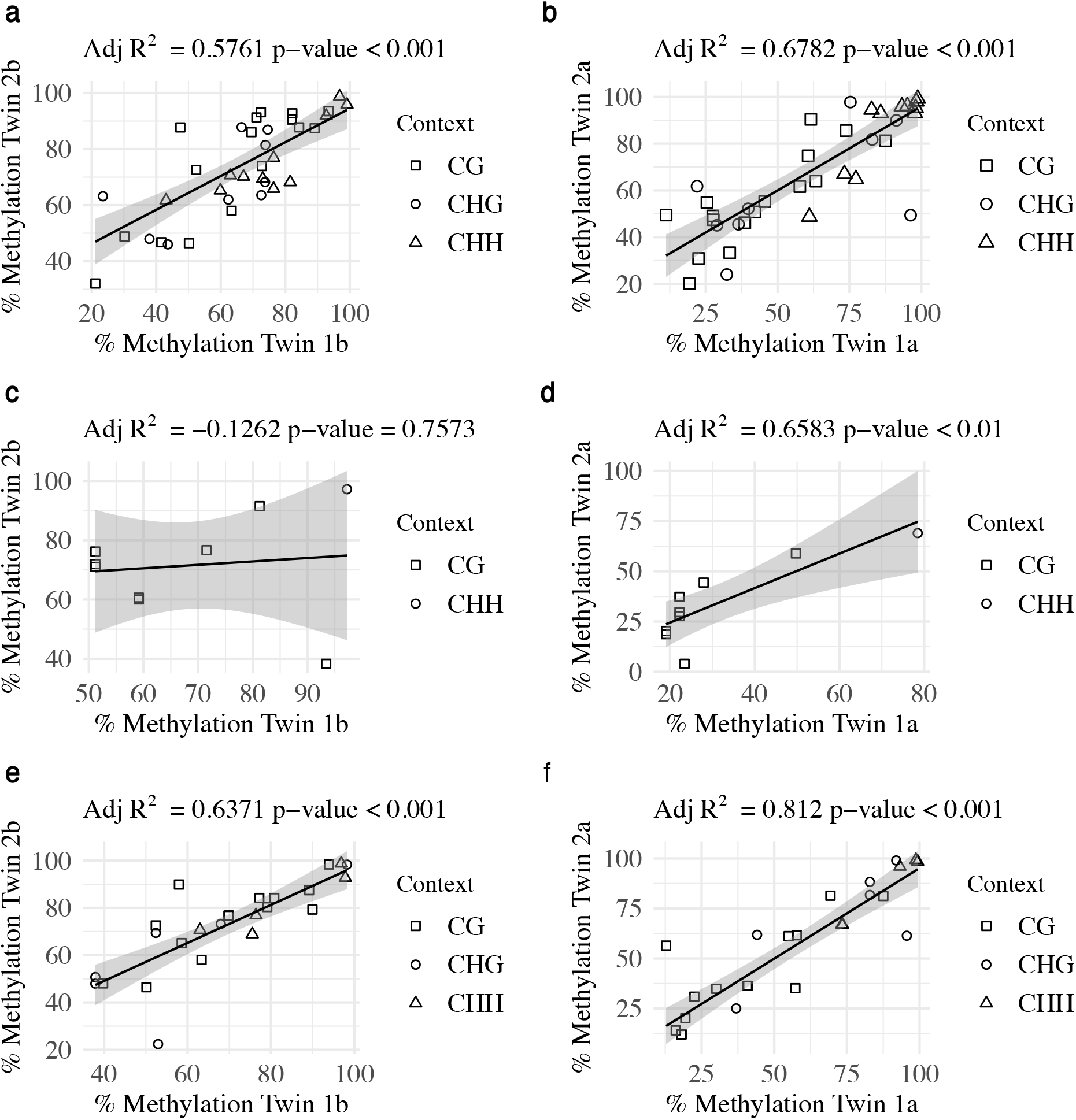
Linear regression of percent methylation within shared regions with significant differential cytosine methylation in no-BF ‘Stukey’ twins (**a, c, e**) and BF ‘Stukey’ twins (**b, d, f**). Panels **a** and **b** represent methylation in DMRs upstream (within 2,000 bp) of a gene, panels **c** and **d** represent methylation in intragenic DMRs, and panels **e** and **f** represent methylation in DMRs downstream (within 2,000 bp) of a gene. Each circle represents the percent methylation in each twin in a single region that is significantly differentially methylated in both ‘Stukey’ twin pairs. Red circles represent DMRs in the CG context, green triangles represent DMRs in the CHG context, and blue squares represent DMRs in the CHH context.

**Fig. S8.**
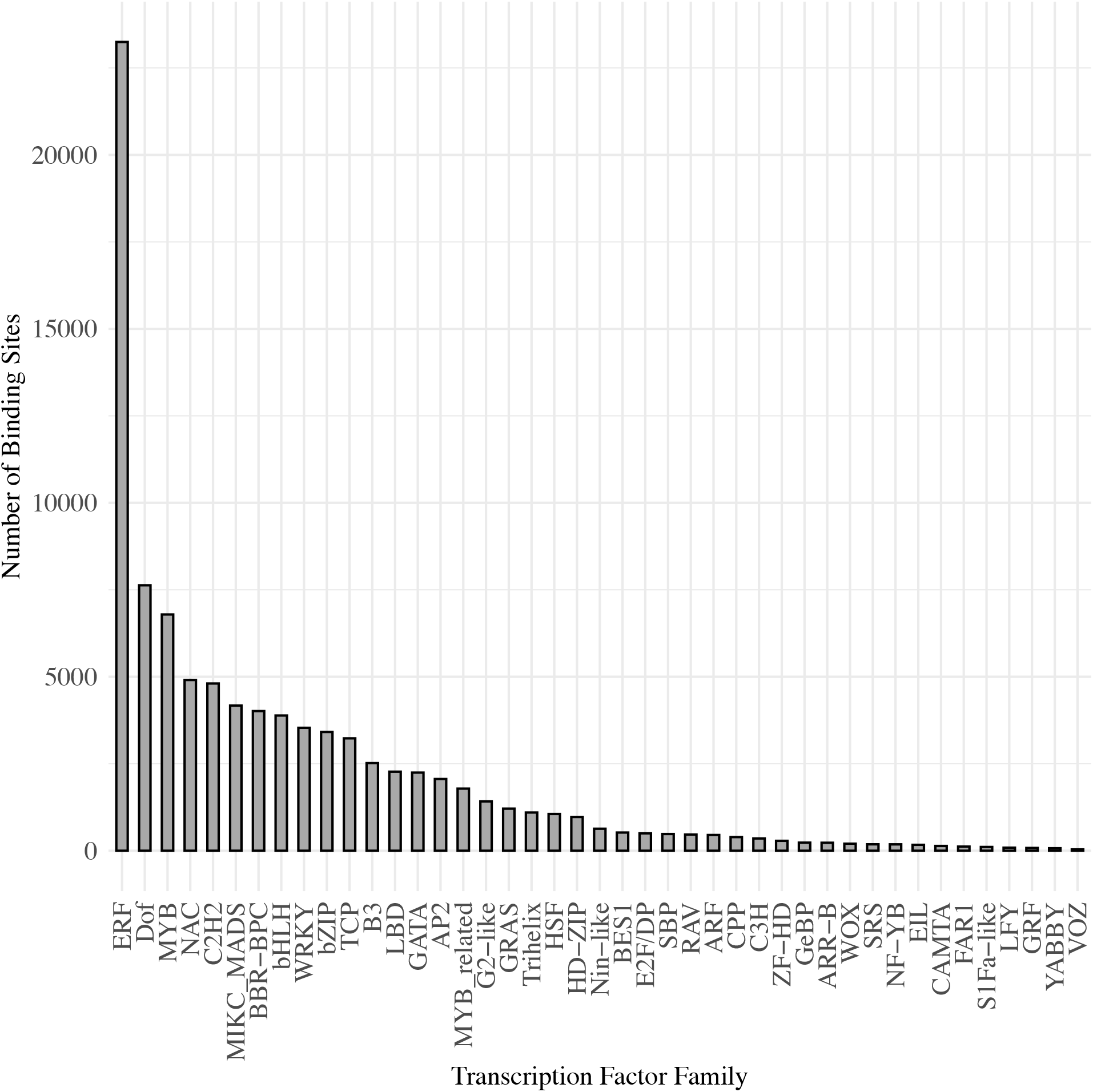
Number of transcription factor binding sites identified in the shared DMRs in all methylation contexts.

**Fig. S9.**
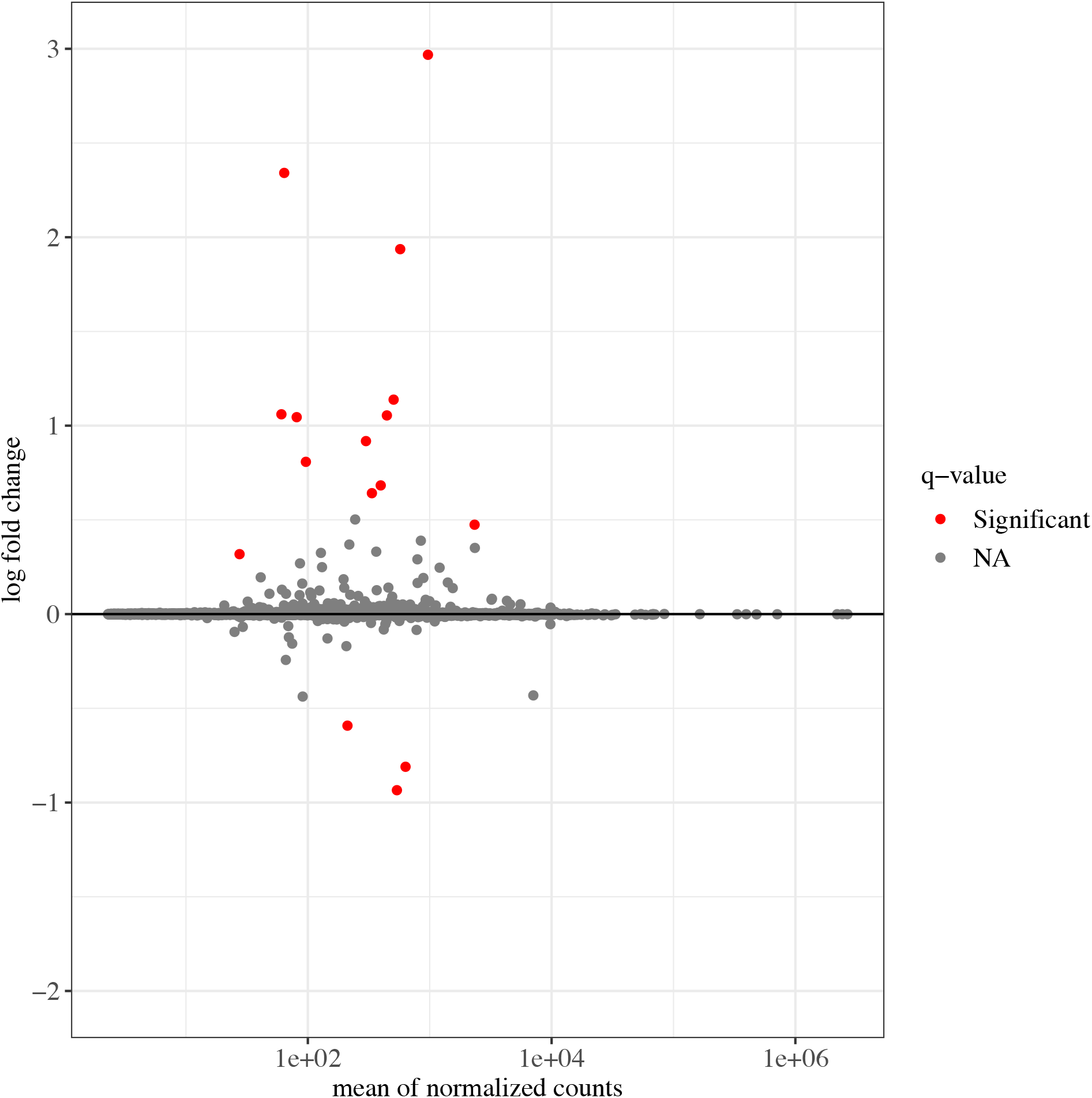
Differentially expressed genes in contrast comparing condition (BF vs no-BF). Red points represent those defined as significantly differentially expressed in each contrast (p-value < 0.1).

**Fig. S10.**
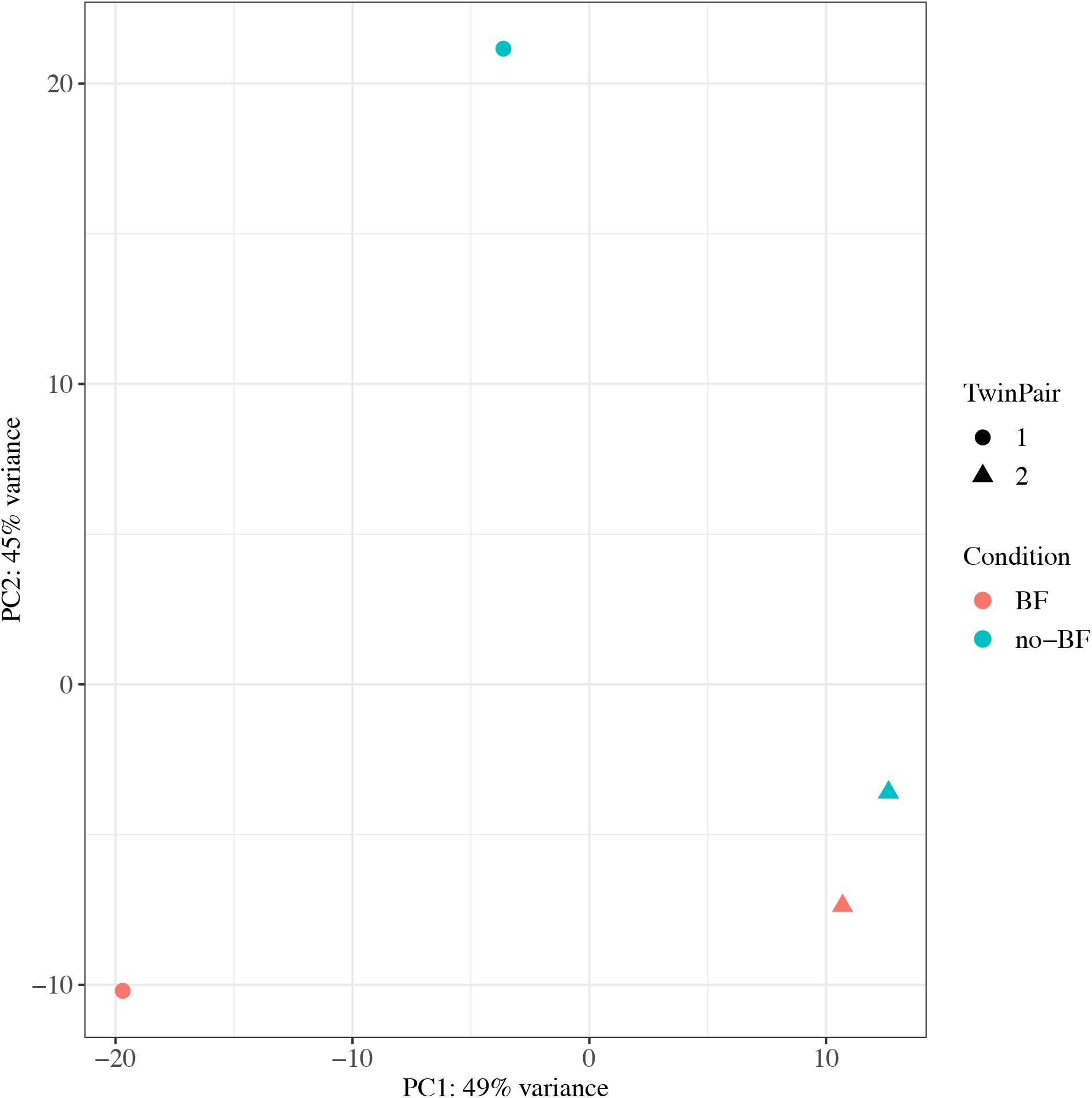
Principal component analysis of differential gene expression data showing separation by ‘Stukey’ twin pair and BF condition.

