## Supplementary Data S2 for "Integrated analysis of the methylome and transcriptome of twin almonds (*Prunus dulcis* [Mill.] D.A.Webb) reveals genomic features associated with non-infectious bud failure"

>Scaffold_6_UnidentifiedProtein

TCATGCAGCTGGAGCATAATCATAACCACCATCATCATCGTCATCGTCATCCCCTTCCATGGCTGCTGGAGCATAGTCAT

AGTCTGAATCATCGTCGTCGCCGTCGCCTTCTATACATGCTGCCAGGGCAAATATATCATCACCGTTGAAGTCGTCGACT

CCATCGCCATGCACCTCGTACCATGAGGCACTGATGATCAACTTGGAAAGGGGAAACAT

>Scaffold_6_UnidentifiedProtein

MFPLSKLIISASWYEVHGDGVDDFNGDDIFALAACIEGDGDDDDSDYDYAPAAMEGDDDDDDDGGYDYAPAA
